# Adaptation to *ex vivo* culture drives human haematopoietic stem cell loss of repopulation capacity in a cell cycle independent manner

**DOI:** 10.1101/2022.11.17.516906

**Authors:** Carys S. Johnson, Kendig Sham, Serena Belluschi, Xiaonan Wang, Winnie Lau, Kerstin B. Kaufmann, Gabriela Krivdova, Emily F. Calderbank, Nicole Mende, Jessica McLeod, Giovanna Mantica, Matthew J. Williams, Charlotte Grey-Wilson, Michael Drakopoulos, Shubhankar Sinha, Evangelia Diamanti, Christina Basford, Anthony R. Green, Nicola K. Wilson, Steven J. Howe, John E. Dick, Bertie Göttgens, Natalie Francis, Elisa Laurenti

## Abstract

Loss of long-term haematopoietic stem cell function (LT-HSC) hampers the success of *ex vivo* HSC gene therapy and expansion procedures, but the kinetics and the mechanisms by which this occurs remain incompletely characterized. Here through time-resolved scRNA-Seq, matched *in vivo* functional analysis and the use of a reversible *in vitro* system of early G_1_ arrest, we define the sequence of transcriptional and functional events occurring during the first *ex vivo* division of human LT-HSCs. We demonstrate that contrary to current assumptions, loss of long-term repopulation capacity during culture is independent of cell cycle progression. Instead it is a rapid event that follows an early period of adaptation to culture, characterised by transient gene expression dynamics and constrained global variability in gene expression. Cell cycle progression however contributes to the establishment of differentiation programmes in culture. Our data have important implications for improving HSC gene therapy and expansion protocols.

## Introduction

A trillion blood cells are produced daily in humans. This impressive output is achieved by rare haematopoietic stem cells (HSCs) with the unique capacity to drive production of all blood cell types (differentiation), while maintaining a functional HSC pool (self-renewal). Owing to this extensive regenerative capacity, HSCs are the key functional units of HSC transplantation, used in clinical setting for more than 40 years, and of HSC gene therapy (GT). HSC GT promises to be the only long-term treatment option for more than 10 monogenic diseases, including immunodeficiencies, hereditary anemias and metabolic disorders (*1*). HSC GT, as well as other potential clinical applications of HSCs, currently require an *ex vivo* culture step, which by yet mostly uncharacterized mechanisms, is associated with a strong reduction in HSC functionality (*2, 3*). Minimising HSC functional attrition during *ex vivo* protocols would provide patients with larger numbers of functional HSCs. This would not only improve treatment efficacy and safety in current HSC GT indications, but also open the way for new therapeutic applications.

Long-term blood formation post-transplantation is driven by highly quiescent long-term HSCs (LT-HSCs) (*4, 5*). Quiescence (G_0_) is defined as the reversible absence of cell cycling and is characterised by decreased cell size, reduced protein biosynthesis (*6, 7*), high autophagic recycling (*8, 9*), an elevated basal expression of cell stress response pathways such as the UPR (*10, 11*) and ERAD (*12, 13*), and a relatively inactive glycolytic metabolism (*14, 15*) maintained *in vivo* by the hypoxic bone marrow niche (*16, 17*). Upon receiving a mitogenic stimulus, HSCs undergo quiescence exit (G_0_ to end of early G_1_) and progressively phosphorylate Retinoblastoma (Rb) until the restriction point (boundary of early – late G_1_), where commitment to division occurs and subsequent cell cycle progression (late G_1_ – S – G_2_– M phases) ensues. CDK6 is a master regulator of HSC quiescence exit kinetics, whose differential regulation within the HSC pool guarantees long-term maintenance of HSC pool size (*18*). An emerging body of work has demonstrated further heterogeneity within the quiescent state of LT-HSCs associated with distinct lineage preferences and kinetics of reconstitution post-transplantation (*19–22*). Current GT protocols target the CD34^+^ fraction (a heterogeneous mix of HSCs and progenitor cells), but there is currently a limited understanding of how these *ex vivo* protocols impact the underlying biology unique to the LT-HSC population.

In current *ex vivo* culture systems, HSCs inevitably exit quiescence and undergo cell cycle progression. *Ex vivo* culture increases protein synthesis rates (*6, 23*), remodels the mitochondria to an activated state (*24*) with increased oxidative metabolism and ROS production (*25*), disrupts optimal proteostasis programs (*23, 26*) and reduces the dependency on lysosomal recycling (*27*). These changes are all associated with a net decline in long-term repopulation capacity. It has been presumed from literature based on *in vivo* models (*28–30*) that cell cycle progression itself drives loss of HSC function in culture. Early studies show that it is exclusively haematopoietic stem and progenitor cells (HSPCs) in the G_0_ phase of the cell cycle before (*31*) and following culture (*32, 33*) that engraft, but not their cycling counterparts. However, to date no study has formally addressed this question in human LT-HSCs undergoing clinically relevant culture. In addition, a number of strategies have been developed to maintain or expand HSCs *ex vivo* (*34*), some of which are extremely promising in both research and clinical settings. Despite these advances, we are still lacking a mechanistic understanding of the interplay between division and maintenance of self-renewal *ex vivo*.

In this study, by pairing scRNA-Seq with *in vivo* functional analysis in a time-resolved manner, we dissect the transcriptional and functional changes occurring in highly purified human LT-HSCs over their first division in culture. Our data globally identify an early time window of adaptation to the culture conditions, during which human LT-HSC transcriptional programmes are deeply remodelled and which precedes and sets the stage for successive loss of HSC function and initiation of differentiation. In addition, by pharmacologically preventing human LT-HSCs from progressing into the late G_1_ phase of the cell cycle, we demonstrate that cell cycle progression does not contribute to the decline in LT-HSC long-term repopulation capacity observed *ex vivo* but facilitates the establishment of lineage priming programmes.

## Results

### Loss of HSC function occurs early during *ex vivo* culture

HSC functional attrition *ex vivo* (*2, 3*) severely hampers the efficacy of clinical protocols reliant on culture prior to transplantation. Loss of HSC function is commonly thought to be driven by quiescence exit and subsequent cell cycle progression, with dynamics that, to our knowledge, have not been reported to date. We therefore sought to study the kinetics of HSC functional attrition using purified human LT-HSCs, choosing early timepoints of culture (≤ 72 h), corresponding to key transitions of cell cycle progression. Here we opted to study two distinct *ex vivo* systems. In the first, we sought to mimic GT conditions and therefore used mobilized peripheral blood (mPB) LT-HSCs (flow sorted as CD34^+^ CD19^-^ CD38^-^ CD45RA^-^ CD90^+^ CD49f^+^ (*4*)) cultured in a 62 h *ex vivo* GT protocol including two hits of transduction with a lentiviral vector (LV) expressing GFP (*2, 35, 36*). In the second, we aimed to examine conditions commonly used in experimental settings (“EXPER” conditions). Cord blood (CB) LT-HSCs, isolated using the same markers above, were cultured for 72 h in differentiation facilitating media promoting cell cycle progression and lineage commitment (*18*). We first characterised the cell cycle kinetics of LT-HSCs in our two distinct *ex vivo* conditions. We used phosphorylation of Rb at the serine residue 807-811 as a marker of the transition through the restriction point from early to late G_1_, determining the time at which quiescence exit is complete (G_0_ – early G_1_) and cells have entered cell cycle progression towards division (late G_1_-S-G_2_-M). LT-HSCs cultured in either GT or EXPER conditions both entered cell cycle progression (late G_1_) at approximately 24 h post culture initiation (EC_50_ of 24.7 h for GT and 23.9 h for EXPER) as assessed by flow cytometry (**Fig 1A-B**). Concurrently, the proportion of cycling cells (S-G_2_-M phases) increased within the 24-72 h window (**Fig.1C-D**) as determined by pRb/DAPI cell cycle analysis. LT-HSC’s time to first division at single cell resolution was estimated at 62.0 h ± 6.5 h for GT conditions and 53.4 h ± 6.5 h for EXPER conditions (**Fig. 1E-G**). Taken together these data demonstrate that independently of HSC source and culture conditions, LT-HSCs complete quiescence exit by approximately 24 h and their first division in culture by 72 h.

**Fig. 1.**
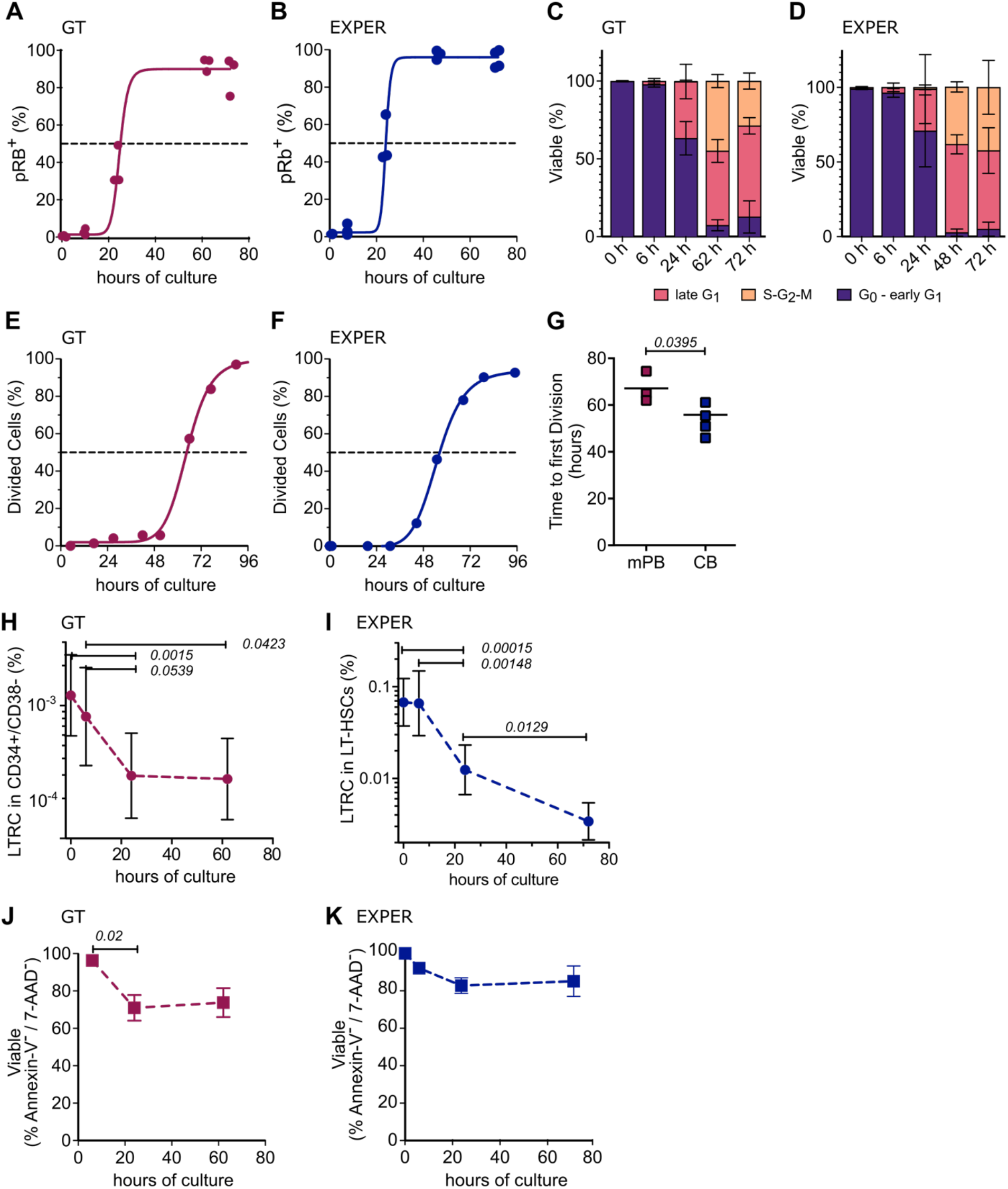
Kinetics of cell cycle progression, survival and loss of long-term repopulation capacity of LT-HSCs during *ex vivo* culture. **(A-B)** Cumulative quiescence exit kinetics of LT-HSCs cultured in GT **(A)** and EXPER **(B)** systems determined by phospho Rb (Ser 807 – 811) flow cytometry analysis. **(A)** n=3 biological replicates for 0 h, 24 h, 62 h, 72 h and n=4 biological replicates for 6 h. **(B)** n=3 biological replicates for 0 h, 6 h, 24 h, 48 h and n=4 biological replicates for 72 h. Curve is least-squares sigmoidal fit with EC_50_ = 24.72 h; Standard error ± 5.186; R^2^ = 0.9797 **(A)** and EC_50_ = 23.94 h; Standard error ± 3.944; R^2^ = 0.9844 **(B)**. Dashed line indicates EC_50_ and determined time of quiescence exit. **(C-D)** Cell cycle phase assignment of LT-HSCs cultured in GT **(C)** and EXPER **(D)** systems determined by pRb/DAPI flow cytometry analysis. Equivalent repeats as in (A) for GT and (B) for EXPER. **(E-F)** Cumulative first division kinetics (excluding dead cells) of LT-HSCs cultured in GT **(E)** and EXPER **(F)** systems. Curve is least squares sigmoidal fit. Representative examples shown (biological replicates: n=3 for GT and n=4 for EXPER. Dashed line indicates EC_50_ and determined time to first division. EC_50_ = 64.83 h; 95% CI = 62.83 h – 66.87 h; R^2^ = 0.9976 **(E)** and EC_50_ = 55.44 h; 95% CI = 54.28 h – 56.70 h; R^2^ = 0.9995 **(F).** **(G)** Time to first division kinetics summary of LT-HSCs cultured in GT **(E)** and EXPER **(F)** systems. (biological replicates: n=3 for GT; n=4 for EXPER). Unpaired t-test shown. **(H-I)** Percent of LTRC in CD34^+^/CD38^-^ cells cultured in GT **(H)** or LT-HSCs cultured in EXPER **(I)** systems as determined by LDA analysis in the transplanted population, and calculated with ELDA statistics. % LTRCs at each time point +/- 95% CI shown, raw data in **Table S1**. GT: 0 h: 29 mice; 6 h: 21 mice; 24 h: 23 mice; 62 h: 23 mice. EXPER: 0 h: 31 mice; 6 h: 19 mice; 24 h: 31 mice; 72 h: 40 mice. Chi-squared test performed between groups and significant values indicated. **(J-K)** Survival of LT-HSCs cultured in GT **(I)** and EXPER **(J)** systems determined by Annexin-V/7-AAD flow cytometry. n=3 biological replicates at each time point. Paired t-test shown for 6 h and 24 h comparison. Mean +/- SD shown.

Next, to quantify how long-term repopulating frequency is affected by *ex vivo* culture, we performed limiting dilution analysis (LDA) xenotransplantation assays with NSG immunocompromised mice. To this end, we transplanted fixed numbers of i) mPB CD34^+^ CD38^-^ cells cultured in GT conditions for 0 h (G_0_), 6 h, 24 h (end of early G_1_), or 62 h (time of infusion into patients in HSC GT protocols); ii) CB LT-HSCs cultured in EXPER conditions for 0 h (G_0_), 6 h, 24 h (end of early G_1_), or 72 h (post first division). Human engraftment was analysed by flow cytometry 18 weeks post-transplantation. Of note, in the GT conditions, almost all animals displayed grafts with GFP^+^ cells (**Data S1**). This broadly indicates that our *in vitro* culture system is robustly recapitulating a GT protocol and that transduced cells contribute to long-term engraftment. LDA showed that the frequency of long-term repopulating cells (% LTRC) within the transplanted LT-HSC population was unchanged for the first 6 h of culture (GT: *p= 0*.*568* ; EXPER: *p=0*.*938*). By 24 h, the time at which LT-HSCs have completed quiescence exit, the % LTRC dropped conspicuously (GT: ∼3 fold, p= *0*.*0539*; EXPER: ∼5 fold, *p=0*.*00148* compared to 6 h). An additional decrease in the % LTRC was observed past 24 h in EXPER conditions (∼2 fold, *p=0*.*0129* compared to 24 h), when LT-HSCs are progressing through the cell cycle and entering mitosis (**Fig.1H-I, Data S1**).To verify if the significant decrease in the % LTRC observed between 6 and 24 h post-culture initiation is due to cell death, we measured apoptosis via Annexin-V/7-AAD staining. A decrease in viable cells was observed between 6 h and 24 h *ex vivo* with no further decrease at later time points in both GT and EXPER culture systems (**Fig.1J-K**). Interestingly, the loss in LTRC observed between 6 h and 24 h *ex vivo* largely outnumbered the loss in viability measured in the same time window. We therefore conclude that HSC loss *ex vivo* occurs predominantly before cells enter the late G_1_ phase of the cell cycle, with kinetics that cannot be primarily explained by cell death.

### LT-HSCs’ transcriptome dynamics over the first cell cycle *ex vivo*

To better understand the kinetics of LTRC loss *ex vivo*, we next investigated the transcriptional changes associated with the first cell division of LT-HSCs in culture. Flow-sorted CB LT-HSCs were cultured for 0 h, 6 h, 24 h and 72 h in EXPER medium before scRNA-seq via an adapted Smart-Seq2 protocol (*37*). 666 cells were sequenced over 2 independent experiments and 429 cells (64%) passed quality control. Cells from both experiments were integrated via 2 distinct pipelines (ScanPy and Seurat 4; see Methods), which yielded highly concordant results. Dimensionality reduction was performed following selection of highly variable genes through the 2 independent packages. *Ex vivo* cultured LT-HSCs group by culture duration in UMAP visualisations, independently of the batch (**Fig. 2A, S1A**), of whether cell cycle regression was applied (**Fig. S1B)** and of the integration pipeline used (**Fig. S1C**).

**Fig. 2.**
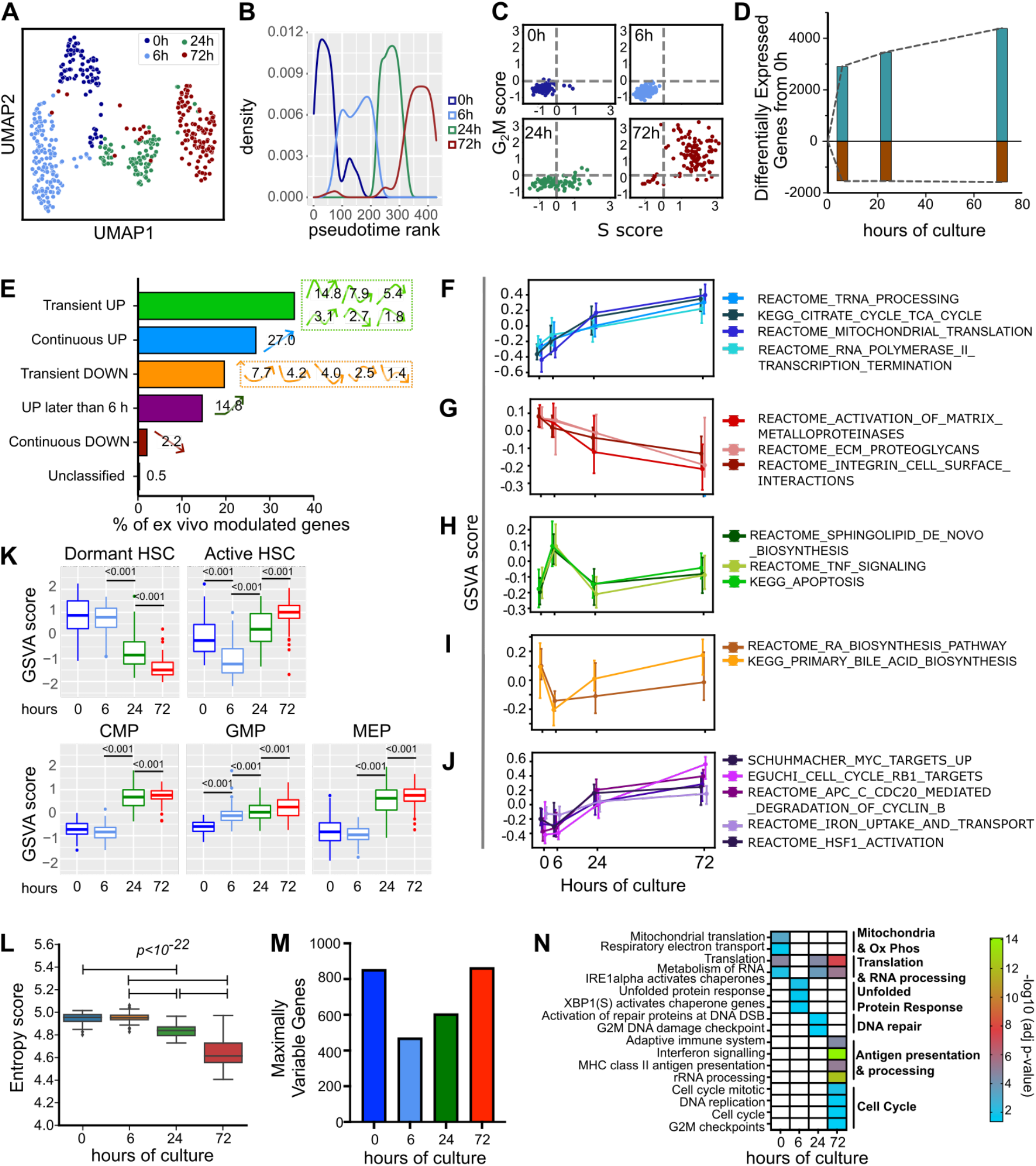
Dynamics of gene expression over the first division of LT-HSC *ex vivo* at single cell resolution. **(A)** UMAP of 429 single LT-HSCs cultured in the EXPER system over a time-course of 0 h, 6 h, 24 h and 72 h (n=2 independent experiments). UMAP generated using Seurat 4 pipeline following cell cycle regression. **(B)** 2D pseudotime rank plot of LT-HSCs over time-course generated following cell cycle regression. **(C)** Transcriptional allocation of cell cycle status at time-points of LT-HSC culture; n= 429 single cells. **(D)** Number of differentially expressed genes (FDR<0.05) at each time-point with respect to 0 h. Full list of genes available in **Data S3.** **(E)** Broad patterns of gene expression identified over time-course (8,966 genes classified after filtering by DEG Report algorithm). Numbers indicate the percentage of genes showing the specific patterns of gene expression displayed to the right of bar. **(F-J)** GSVA score of c2 curated pathways showing specific expression patterns: **(F)** continuous up; **(G)** continuous down; **(H)** transient up; **(I)** transient down; **(J)** up later than 6h. GSVA score calculated per single cell with line at median and upper and lower whiskers indicate 25th and 75th percentile of expression. **(K)** GSVA scores of indicated published gene signatures, representative of specific HSPC subsets. Median and interquantile range shown. **(L)**scEntropy value at each timepoint (calculated for both batches combined; Wilcoxon rank sum test performed and significant values indicated; 0 h vs 6 h p = 0.835). Median and interquantile range shown. **(M)** Number of maximally variable genes at each timepoint (MVG; see methods; 2792 genes total). **(N)** Selected biological pathways significantly enriched from MVG (-log_10_(adjusted p-value) <0.05). Full list of pathways available in **Data S6**.

An increasing body of work is identifying transcriptional and functional heterogeneity within the human LT-HSC pool (*19–22, 38*), indicating the potential presence of multiple subpopulations at the 0 h time-point. 0 h LT-HSCs were therefore further split into 2 clusters by k means clustering (**Fig. S1D)**. Differential gene expression was performed with DESeq2, followed by GSEA and GSVA analyses (**Data S2)**. One cluster expressed significantly higher levels of genes involved in hypoxia regulation (**Fig.S1E**) than the other and displayed enrichment for the most dormant (*22*) and multipotent (*19*) LT-HSCs gene signatures (**Fig.S1F)**. This cluster was denoted “0 h-early” and was subsequently used as pseudotime origin (**Fig.S1G**). In the other cluster, named “0 h-late”, we found higher levels of oxidative phosphorylation genes than in “0 h-early” LT-HSCs (**Fig.S1H;** *p=0*.*0580*) and significant enrichment of gene signatures related to a myelo-lymphoid restricted self-renewing HSC subset (*19*) (**Fig.S1F**). In summary, our scRNA-seq strategy could identify transcriptional heterogeneity within the quiescent human LT-HSC fraction, in line with recent literature (*19–22, 38*).

Following the 0 h time-point, pseudotime ordering largely recapitulated chronological time (**Fig. 2B, Fig. S1I**) and in agreement with functional analysis, transcriptional assignment of cell cycle status showed an increasing proportion of LT-HSCs allocated to S-G2-M phases as culture progresses (**Fig. 2C**). LT-HSCs therefore progress through distinct transcriptional states over time in culture. To better determine transcriptional kinetics over the 72 h time-course, we first performed differential gene expression for all pairwise comparisons in the dataset using DESeq2 (**Data S3**). *Ex vivo* culture has an extensive and rapid effect on the transcriptome with 10,010 genes changing over the time-course (all genes differentially expressed in at least one pairwise comparison, hereafter termed “*ex vivo* modulated genes”, **Data S4)**. Interestingly, 75% of gene expression change from 0 h to 72 h is observed within the first 6 h of culture (4460 / 5980 genes) (**Fig. 3D**), before the vast majority (>90%) of cells have undergone quiescence exit in these culture conditions (**Fig.1A**). We then used the degPatterns package (*39*) on *ex vivo* modulated genes to define gene expression patterns over the time course (**Fig. 2E, Fig. S2A, Data S5**). In parallel, Gene Set Variation Analysis (GSVA) (*40*) was performed on the *ex vivo* modulated genes to identify biological pathways changed during culture, and pathways were again clustered by DEG Report (**Fig. S2B, Data S5**). Similar dynamic patterns of upregulation and downregulation were observed at the gene and pathway level (**Fig.S2A-B**).

**Fig. 3.**
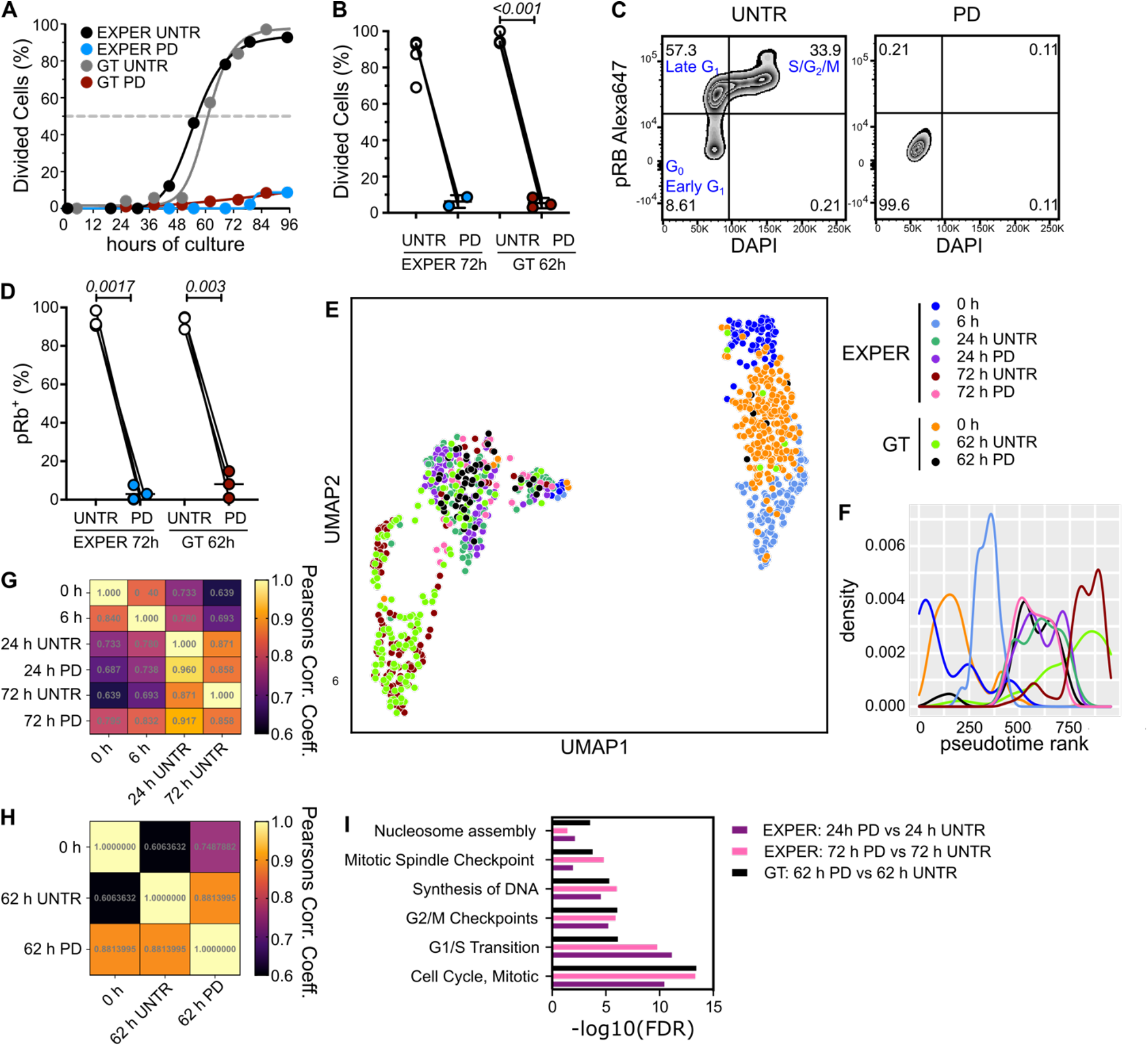
Transcriptional effects of preventing progression past early G_1_ during *ex vivo* culture of LT-HSCs. **(A)** Cumulative first division kinetics (excluding dead cells) of UNTR/PD treated LT-HSCs cultured in GT (grey and dark red) or EXPER (black and blue) system. Curve is least squares sigmoidal fit. Representative example shown (n=3 UNTR/PD matched biological replicates for GT and n=2 UNTR/PD treated matched biological replicates for EXPER). Dashed line indicates EC_50_ and determined time to first division. **(B)** Divided single cells as a proportion of total alive cells at 96 h (EXPER: n=5 biological repeats, n=2 UNTR/PD matched biological repeats; GT: n=3 matched biological repeats). Paired t-test. **(C)** Representative example of flow cytometry plot for phospho Rb (Ser 807 – 811) Alexa 647 and DAPI staining on UNTR (left) or PD treated (right) CB LT-HSCs cultured in EXPER medium for 72 h. **(D)** Quantification of pRb+ (as % of viable cells) in UNTR/PD treated LT-HSCs cultured for 62 h in GT (n=3 UNTR/PD treated matched biological repeats; no LV transduction) or 72 h in EXPER (n=3 UNTR/PD treated matched biological repeats) systems. Paired t-test. **(E)** UMAP visualisation of scRNAseq from 954 LT-HSCs from the indicated culture conditions (EXPER: 536 single cells: GT: 418 single cells). Cell cycle regression applied. **(F)** 2D Pseudotime density rank plot of single cells shown in (E). Cell cycle regression applied. **(G)** Pearsons correlation coefficient estimate comparing the median expression value of 10,903 genes at time-point/condition comparisons in the EXPER dataset. 10,903 genes represent the union of all differentially expressed genes between any 2 UNTR timepoints and between PD treated and UNTR conditions used for analysis (available in **Data S4**). **(H)** Pearsons correlation coefficient estimate comparing the median expression value of 5,469 genes at time-point/condition comparisons in the GT dataset. 5,469 genes represent the union of all differentially expressed genes between any 2 UNTR timepoints and between PD treated and UNTR conditions used for analysis (available in **Data S4**). **(I)** Selected Reactome pathways (FDR<0.05) enriched between PD treated and UNTR LT-HSCs cultured for matched durations of 24 h in EXPER (purple), 72 h in EXPER (pink) and 62 h in GT (black) systems. Full DeSeq2 results and Reactome pathway enrichment available in **Data S7**.

GSVA scores of pathways linked to transcription, RNA splicing, translation and oxidative phosphorylation related processes increased cumulatively with time in culture (**Fig.2F**). Conversely, GSVA scores of pathways related to integrin signalling, extracellular matrix (ECM) generation and matrix metalloproteinase activation which are important for HSC *in vivo* niche maintenance (*41*) were continuously downregulated (**Fig. 2G**). This suggests a gradual remodelling of HSC cell adhesion properties upon *ex vivo* culture. In contrast, the AP-1 transcription factors JUN and FOS (**Fig. S2C**) dropped dramatically by 6 h, pinpointing early loss of the quiescent network.

We also observed a number of transient patterns of expression over the first 72 h of culture. Particularly abundant were genes/pathways which show upregulation from 0 h to 6 h and are then decreased at 24 h (27.3 % of total genes; % 76.3 of “Transient UP genes”). GSVA scores of transiently upregulated pathways include sphingolipid *de novo* biosynthesis, TNF signalling and apoptosis (**Fig.2H**). Genes with maximum levels of expression at the 6 h timepoint included key cell stress and apoptotic regulators such as the dual regulators of apoptosis *CFLAR* and *BIRC2*, the master regulator of the unfolded protein response *ATF4* and the ceramide biosynthesis enzyme *DEGS1* (**Fig.S2D**). We also observe transient downregulation of retinoic acid and bile acid biosynthesis pathways (**Fig. 2I**). Collectively, these transient patterns of expression early in the time-course suggest that HSCs initiate a cell stress response to adapt to the instructive signals of culture.

Another expression pattern of interest, which encompasses 14.8% of all genes (**Fig.2E**), is marked upregulation of expression after the 6 h timepoint (**Fig.2K**). MYC targets, cell cycle progression, iron uptake and activation of HSF1 pathway genes (**Fig.2J**) follow this pattern in addition to *MYC* gene expression (**Fig.S2E**). Correspondingly, gene expression signatures associated with active human LT-HSCs (*22*) were found to be upregulated from 6 h to 24 h and further at 72 h (**Fig.2K**). Gene expression signatures of dormant human LT-HSCs exhibited the opposite pattern, decreasing sharply from 6 to 24 h. Differentiation signatures for GMP, CMP and MEP generated from a landscape of CB differentiation (*42*) also showed a “UP later than 6 h” expression dynamic (**Fig.2K**). To quantitatively measure the degree of differentiation in an unbiased manner, we calculated single cell entropy scores using scEntropy (*43*), a parameter which decreases during differentiation. scEntropy scores significantly decreased only after 6 h (**Fig.2L**), supporting the concept that cell fate decisions occur at the 6 – 24 h transition. These data suggest that lineage priming transcriptional programs begin to be activated later than 6 h post culture initiation, concurrently with loss of long-term repopulation capacity.

Dynamic changes in gene expression variability of key gene networks have been linked to cell fate decisions in haematopoietic progenitors (*44*) and T cell activation (*45*). To better understand LT-HSC state transitions *ex vivo*, we therefore investigated the variability of single cell gene expression over the time-course. The BASiCS method was employed (*46–48*) to calculate the overdispersion measure of variability. Differentially variable genes between every pair-wise comparison of timepoints were extracted and genes with maximum variability at each time-point (MVG) were determined (see Methods). The number of MVGs was lowest at 6 h (**Fig.2M**) indicating that gene expression variability is restrained directly preceding loss of HSC function, and then increases again as cells progress through the cell cycle. MVGs displayed expression patterns found in all groups described above in variable proportions (**Fig.S2F**). MVGs at 6 h were enriched for genes involved in UPR regulation (including *DNAJC3* and *XBP1*). At later time-points, MVGs are linked first to translation regulation (at 24 h) then to antigen processing (including *CD74* and *HLA-DRB5*) and cell cycle progression at 72 h (**Fig.2N, Data S6**).

Overall our time-resolved transcriptomic analysis identifies two distinct phases of LT-HSC activation *ex vivo*: i) an early adaptation phase (0 – 6 h), marked by overall restriction in gene variability but transient expression and increased variability in genes associated with cell stress responses; ii) a later phase (post 6 h) marked by MYC activation, expression of genes related to cell cycle progression and initiation of differentiation programmes.

### LT-HSC transcriptomic adaptation to *ex vivo* culture is largely independent of cell cycle progression

During *ex vivo* culture, changes to HSC metabolism and organelle biology are accompanied by progression into late G_1_ (and later cell cycle phases) with loss of HSC function (*23–25, 27, 49*). However, whether cell cycle progression is by itself causative of the loss of HSC function has not been formally examined to date. Our observation that the sharpest drop in LT-HSC repopulation capacity occurs *ex vivo* before most HSCs enter late G_1_ prompted us to formally test to what extent progression past early G_1_ contributes to HSC identity and function in culture.

To achieve this we took advantage of the CDK4/CDK6 inhibitor Palbociclib (PD033299; herein PD), previously established to prevent division of CB LT-HSCs (*18*). First, we repeated this finding with LT-HSCs cultured in both the GT and EXPER culture systems (**Fig.3A-B**). Then, we used phospho-Rb flow cytometry staining to determine that PD treatment arrests CB LT-HSCs (**Fig.3C-D**) and mPB LT-HSCs (**Fig.S3A**) in early G_1_. PD treated LT-HSCs cultured for 72 h and subsequently plated in a single cell differentiation assay in the absence of PD (**Fig.S3B**), generated colonies with a similar efficiency (**Fig.S3C**) and of comparable size (**Fig.S3D**) and lineage (**Fig.S3E**) when compared to untreated (UNTR) cells. These data indicate that PD treatment reversibly arrests LT-HSCs in early G_1_ *ex vivo* without compromising their function and can therefore be used to assess the effect of culture in the absence of cell cycle progression.

To disentangle which transcriptional differences observed in culture depend on cell cycle progression, scRNA-Seq was performed on LT-HSCs treated with PD for 24 h (equivalent to quiescence exit completion) and 72 h (equivalent to mitosis) in the EXPER system as well as LT-HSCs treated with PD for 62 h in the GT system. These data were integrated with 0 h and 6 h cells in a unique embedding via the Seurat4 pipeline for a total of 954 cells (n=6 independent experiments). In line with functional data, LT-HSCs treated with PD were confirmed to be allocated to the G_1_ phase of the cell cycle based on their transcriptome independently of the culture system used (**Fig.S3F-G**). Cultured LT-HSCs pharmacologically arrested in G_1_ localise with 24 h UNTR LT-HSCs on the UMAP, independently of the cell source, culture system, duration of PD treatment (**Fig.3E**) and of whether cell cycle regression was applied or not (**Fig.S3H**). This transcriptional similarity between pharmacologically early G_1_ arrested cells (PD treated) and UNTR LT-HSCs cultured for 24 h is also observed on pseudotime analysis (**Fig.3F, Fig.S3I**). Taken together, these data suggest that transcriptional changes up to 24 h *ex vivo* are largely independent of progression to late G_1_ but those beyond 24 h are linked to cell cycle progression.

We next measured the extent of correlation in gene expression between UNTR and PD treated cells at each of the time points. Considering *ex vivo* modulated genes plus genes which are changed as a result of PD treatment (**Data S4**), in the EXPER system, the Pearson’s correlation co-efficient was lowest for 0 h compared to 72 h UNTR cells as expected (**Fig.3G**) and was the highest for PD treated cells cultured for matched durations (24 h UNTR vs 24 h PD = 0.960; 72 h UNTR vs 72 h PD = 0.858) (**Fig.3G**). Similar results were obtained for the GT culture system (**Fig.3H**). This indicates that most gene expression changes are driven by the amount of time in culture, whereas pharmacological early G_1_ arrest has a minimal global effect on gene expression. Reactome pathway analysis demonstrated that the majority of pathways significantly enriched in genes changed between PD and UNTR cells cultured for matched durations relate to processes involved in cell cycle progression (**Fig.3I, Data S7**). These data demonstrate that PD treatment in both culture systems predominantly impacts expression of genes linked to cell cycle progression with relatively few off-target effects. We conclude that *ex vivo* culture rapidly and extensively rewires the LT-HSC transcriptome and that most transcriptional changes acquired during 72 h of culture are largely independent of cell cycle progression past early G_1_.

### Loss of long-term repopulation capacity *ex vivo* is independent of cell cycle progression

Next we tested whether inhibition of cell cycle progression past early G_1_ *ex vivo* impacted HSC functional hallmarks. An increase in cell size (**Fig.4A**) and mitochondrial activity (**Fig.4B**), both hallmarks of HSC activation (*15, 25, 49*) were observed over LT-HSC culture in EXPER conditions but were unchanged upon pharmacological G_1_ arrest by PD, demonstrating that cell growth and mitochondrial metabolism are independent of cell cycle progression.

**Fig. 4.**
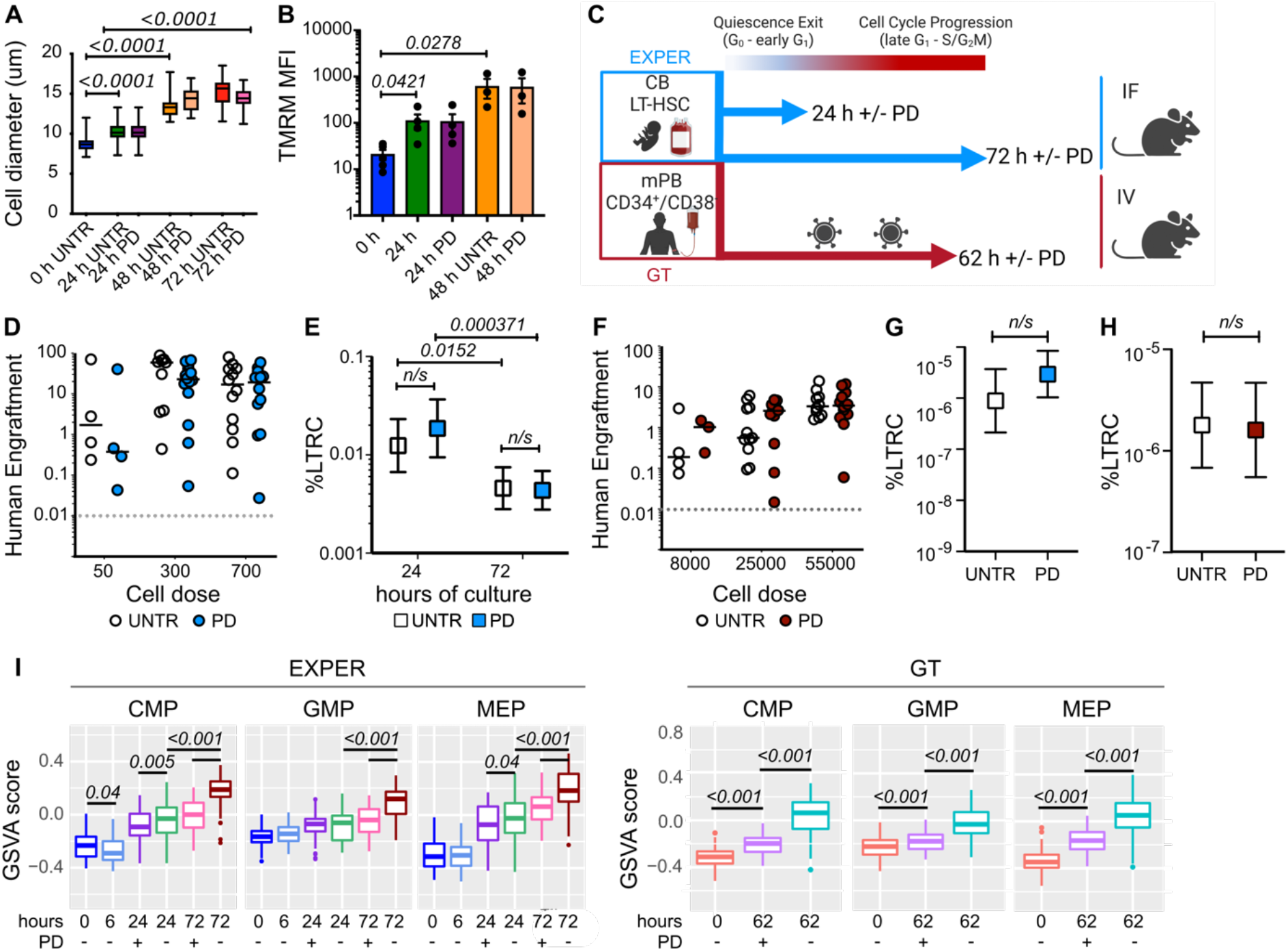
Preventing progression past early G_1_ during *ex vivo* culture of LT-HSCs does not affect loss of long-term repopulation capacity but dampens the establishment of differentiation programmes. **(A)** Cell diameters of single CB LT-HSCs cultured in EXPER conditions (n=2 experiments representing n=469 total single cells). Unpaired t-test. **(B)** Tetramethylrhodamine (TMRM) staining of bulk CB LT-HSCs cultured in EXPER conditions (n=5 biological repeats for 0 h; n = 4 matched biological repeats for UNTR/PD treated 24 h; n=3 matched biological repeats for UNTR/PD treated 48 h). Unpaired t-test. **(C)** Workflow of *in vivo* transplantation of UNTR/PD treated CB LT-HSCs cultured in EXPER conditions and mPB CD34^+^/CD38^-^ cells culture in GT system. Created with BioRender. **(D)** Graft size (% of human CD45^++^ and GlyA^+^) at 18 week post transplantation of UNTR/PD treated LT-HSCs cultured for 72 h in EXPER system (n=5 biological experiments; n=42 PD mice, n = 38 UNTR mice). **(E)** % LTRC in LT-HSCs cultured in EXPER system in presence or absence of PD, determined at 24 h (n=31 mice UNTR, n=30 mice PD) and 72 h (n= 42 mice PD, n=38 mice UNTR). Numerical estimates available in **Data S1**. Chi squared test (24h UNTR compared to 24h PD p=0.405; 72 h UNTR compared to PD p=0.878). **(F)** Graft size (% of human CD45^++^ and GlyA^+^) at 18 week post transplantation of mPB CD34^+^CD38^-^ cells after GT protocol culture for 62 h including LV (n=3 biological repeats; n=25 mice UNTR, n=26 mice PD). Mann Whitney test >0.05 for comparison of UNTR and PD at all doses. **(G)** LDA of secondary transplantation experiment from UNTR/PD LT-HSC EXPER primary mice cohort (D-E). Secondary animals were transplanted with sorted CB CD45^++^ from primary recipients (n= 20 mice total; 10 UNTR, 10 PD; n=1 experiment). UNTR estimate = 1/1129479 (Lower = 1/4712177; Upper = 1/270729) PD estimate = 1/339688 (Lower = 1/967794; Upper = 1/119228). Chi-squared test performed (p=0.190). **(H)** LDA of secondary transplantation experiment from UNTR/PD LT-HSC GT primary mice cohort **(F)**. Secondary animals were transplanted with whole mouse BM isolated from primary recipients (n=21 mice total; UNTR = 11 mice; PD = 10 mice; n=1 experiment). UNTR estimate = 1/557781 (Lower = 1/1465425; Upper = 1/212307). PD estimate = 1/622437 (Lower = 1/1815440; Upper = 1/213407). Chi-Squared test performed (p= 0.860). **(I)** GSVA scores of indicated lineage gene expression signatures from (*42*) at indicated time-points of LT-HSC culture. GSVA score generated per cell and line at median. EXPER: n= 536 single cells; GT: 418 single cells.

To formally determine if progression past early G_1_ contributes to loss of long-term repopulation capacity in the culture conditions tested, we transplanted into NSG mice: i) limiting doses of CB LT-HSCs cultured in presence or absence of PD in EXPER conditions for either 24 or 72 h; ii) mPB CD34^+^/CD38^-^ cells cultured in the presence or absence of PD in GT conditions (**Fig.4C**). The BM from the animals was then analysed for human engraftment after 18 weeks (**Data S1, Fig.S4A**). We observed similar graft sizes and lineage outputs in mice transplanted with CB LT-HSCs cultured in presence or absence of PD for either 24 h (**Fig. S4B-C**) or 72 h (**Fig.4D, Fig.S4D**). Limiting dilution analysis (LDA) at both timepoints of the EXPER LT-HSC culture found no difference in the % LTRC observed both in PD treated and UNTR cultured cells at 24 h (*p=0*.*405*) and 72 h (*p=0*.*878*) **(Fig.4E. Data S1**). This indicates that cell cycle progression does not drive loss of long-term repopulation capacity in under EXPER culture conditions. In fact, the % LTRC for LT-HSCs treated with PD for 72 h is significantly lower (approximately 4 fold; *p=0*.*000371*) than that of LT-HSCs treated with PD for 24 h (**Fig.4E**), further supporting this conclusion. Similarly, in GT conditions, mPB CD34^+^/CD38^-^ pharmacologically arrested in early G_1_ for 62 h show no difference in graft size (**Fig.4F**) and lineage output (**Fig.S4E**) to those of UNTR control animals. Secondary transplantation experiments revealed that the frequency of serially-transplantable HSCs in the injected population was comparable between PD and UNTR conditions for both EXPER (**Fig.4G**; *p=0*.*190*) and GT (**Fig.4H**; *p=0*.*860*) conditions with no significant difference in graft size observed (**Fig.S4F-G**). Taken together, our transplantation experiments demonstrate that long-term repopulation capacity significantly drops over culture, and that cell cycle progression past early G_1_ is not the main driver of such loss of human LT-HSC function.

### Cell cycle dependent initiation of differentiation in cultured LT-HSCs

Cell fate decisions in individual self-renewing HSCs drive lineage specification and are associated with upregulation of lineage specific markers. To understand if progression past early G_1_ contributes to initiation of differentiation, the expression of gene signatures specific to myelo-erythroid progenitors was compared between cultured PD treated and UNTR single LT-HSCs. Compared to UNTR conditions, MEP, CMP and GMP signatures were all significantly downregulated in LT-HSCs treated with PD in either EXPER (72 h) or GT conditions (62 h, **Fig.4I**). Notably, LT-HSCs treated with PD in EXPER conditions expressed significantly less of the characteristic neutrophil granule protease genes *PRTN3, ELANE* and *CTSG* (*50*) (**Fig.S4H**) and of genes linked to erythropoiesis *TPI1*(*51*) (**Data S7**). These data demonstrate that overall, lineage specification is diminished or delayed in absence cell cycle progression suggesting that progression through S-G_2_-M facilitates the transcriptional onset of differentiation.

## Discussion

E*x vivo* culture is known to affect the function and molecular circuits of human HSPCs. Our study extends these findings to highly purified human LT-HSCs, the cells uniquely responsible for long-term outcomes of clinical HSC transplantation, and dissects the sequence of transcriptional and functional events occurring during LT-HSC first *ex vivo* division (**Fig.S4I**). In addition, we conclusively demonstrate that progression through late G_1_ – S – G_2_ and M phases of the cell cycle does not cause the drastic loss in long-term repopulation capacity observed over the first division in culture, but contributes to robust establishment of differentiation programmes.

Our analysis establishes that a profound transcriptional rewiring of the quiescent LT-HSC state occurs very early on *ex vivo*, already within the first 6 hours of culture. While long-term repopulation capacity is fully preserved at this stage, these early gene expression dynamics reflect the drastic change in environmental factors that the cells are exposed to and set the stage for later loss of HSC function. Interestingly, the subsets of quiescent LT-HSCs with distinct transcriptional cell cycle priming and oxidative metabolism observed at 0 h, and previously reported by other studies (*19, 20, 22, 52*) were no longer readily identifiable at 6 h of culture. In addition, the number of maximally variable genes, an indirect measure of intercellular heterogeneity, was lowest at 6 h post culture initiation. We conclude that all LT-HSC subsets adapt to the culture conditions similarly.

Globally our data support a model in which adaptation to *ex vivo* culture is followed by a cell fate decision, whereby LT-HSCs either undergo apoptosis, lose long-term repopulation capacity or rarely, maintain their LT-HSC function. This occurs around the time at which LT-HSCs phosphorylate Rb and enter late G_1_, here estimated between 6 and 24 h post culture initiation in two independent culture settings. Several observations support LT-HSC fate decisions occurring in this time window. First, scEntropy scores, a measure previously reported to track the multiplicity of cell fate potentials (*43, 53*), significantly decreased between 6 and 24 h. Second, in this same timeframe, we observed transient dynamics of expression and/or high intercellular variability in gene sets linked to both HSC survival (ATF4, BIRC2, CFLAR genes and TNF signalling (*54*)) and HSC culling (UPR). High basal levels of stress response pathway expression sets superior “quality control” thresholds for LT-HSCs, resulting in preferential apoptosis in response to oxidative, genotoxic and proteostatic damage (*10–13, 26, 55, 56*). This “culling” of unfit HSCs ultimately maintains the integrity and longevity of the HSC pool *in vivo*. We propose that a similar phenomenon of “HSC quality control” is at play specifically around the time of transition from early to late G_1_ likely determining the balance between survival and differentiation *ex vivo*. Third, transient upregulation of the ceramide biosynthesis enzyme, DEGS1 during the adaptation phase likely contributes to later LT-HSC functional loss. DEGS1 upregulation disrupts lipid homeostasis and underlying proteostatic programmes, with pharmacological inhibition of DEGS1 by 4HPR restoring a coordinated cell stress response program and autophagic recycling to preserve LT-HSC capacity *ex vivo* (*26*). Finally, genes of the retinoic acid signalling pathway, key to maintain HSCs functional features *in vivo* (*29, 57*), are transiently downregulated between 6 and 24 h in culture. Collectively, we conclude that complex and heterogeneous gene expression dynamics unfolding in each single LT-HSC during the adaptation phase will determine whether they maintain or lose function *ex vivo*. Future work will have to experimentally verify how specific adaptation pathways influence HSC function *ex vivo*. In addition, in the clinical setting to minimize the time to engraftment, it is HSPCs (CD34^+^) that are reinfused into patients. Therefore it will be interesting to test if the presence of progenitors might influence the dynamics observed here and/or aggravate functional attrition.

It has long been presumed that LT-HSC division causes loss of LT-HSC function *ex vivo*. Here our study resolves this debate by using a unique experimental system which efficiently and reversibly arrests LT-HSCs progression past early G_1_ *ex vivo*. We conclusively establish that cell cycle progression and division do not drive the sizeable loss of HSC long-term repopulation capacity observed *ex vivo* over 62 -72 h. Importantly, our data do not contradict the large body of previous literature aiming to understand the complex relationship between cell cycle progression and HSC self-renewal capacity. A strong correlation between excessive proliferation of LT-HSCs and eventual loss of self-renewal has been observed through an extensive body of work using *in vivo* mouse models (reviewed in (*58*)). In addition, self-renewal capacity is much enriched in HSCs residing *in vivo* in the G_0_ phase of cell cycle compared to those in G_1_ or S-G_2_-M (*58, 59*). In addition, *ex vivo* studies performed with an heterogeneous population of human HSPCs, demonstrated that the G_0_/G_1_ fraction of cultured CD34^+^ contains the highest frequency of repopulating cells (*33, 60*). Our results complement these findings, by showing that preventing progression of LT-HSCs into the late G_1_-S-G_2_-M phases of the cell cycle is not sufficient to maintain long-term repopulation capacity *ex vivo*.

Differentiation is tightly connected to cell cycle events across many types of stem cells. The duration of G_1_ determines lineage commitment in embryonic (*61–63*) and neural stem cells (*64*). Whether that is the case in HSCs is still under debate (*58, 65, 66*). However, cell cycle regulators such as CDKs and Cyclins are known to control the activity of key regulators of HSC fate either by directly phosphorylating them or by affecting their levels of transcription through chromatin binding (*67–70*). Interestingly, we observe that *ex vivo* upregulation of myelo-erythroid lineage specification gene signatures is dampened when mPB and CB LT-HSCs are prevented from progressing through late G_1_-S-G_2_-M. This indicates that cell cycle progression is not strictly necessary for but facilitates the establishment of differentiative programmes in HSCs. Self-renewal and lineage commitment activities are controlled independently in HSCs both in *in vivo* and *in vitro* contexts in mice (*71–73*). Our study further supports this concept by temporally uncoupling loss of repopulation capacity (occurring before Rb phosphorylation) from establishment of differentiation (occurring post Rb phosphorylation in a partially cell cycle dependent manner) in cultured human HSCs.

The availability of large numbers of functionally fit HSCs remains a major limiting factor for most HSC transplantation and HSC GT patients. The findings of our study have direct relevance for improving current HSC GT protocols and HSC *ex vivo* expansion methods and overcome this barrier. Much optimization to date has been successful at delivering the best gene correction rates within the first 60-70 hours in HSC GT (*1*), and expand the number of functional HSCs present after at least one week of culture (*74, 75*). Our work however pinpoints irreversible HSC fate decisions to the first 24 h of culture, and therefore identifies an untapped window of opportunity to preserve HSC function in any application that requires an *ex vivo* step. We warrant that shortening GT protocols including earlier transductions steps may be useful (*76*), but that should be complemented with approaches aimed at minimizing the loss of HSC functionality occurring early on. Similarly, HSC *ex vivo* expansion strategies may suffer from early bottlenecks that reduce the clonal diversity of HSCs to be expanded, likely presenting long-term risks to patients. We foresee that such bottlenecks could be mitigated by targeting early HSC adaptation responses. In conclusion, this study presents a rich resource that paves the way for novel approaches to deliver significantly increased numbers of functional HSCs to patients.

## Materials and Methods

### Experimental Design

The objectives of this study were to characterise how specific functional and transcriptomic features are affected by *ex vivo* culture. We used two complementary culture systems: 1) EXPER culture conditions: StemPro base media (Thermo Fisher Scientific) supplemented with Nutrients (0.028%) (Thermo Fisher Scientific), Pen/Strep (1%), L-Glu (1%), human LDL (50ng/ml) (Stem Cell Technologies) in addition to the following cytokines: SCF (100ng/ml), Flt-3L (20ng/ml), TPO (100ng/ml), EPO (3 units/ml), IL-6 (50ng/ml), IL-3 (10ng/ml) and GM-CSF (20ng/ml). All cytokines from Miltenyi Biotech except EPO (Janssen). 2) GT culture conditions: GMP SCGM (CellGenix) supplemented with L-Glutamine (1%) (Thermo Fisher Scientific), Pen/Strep (1%) (Thermo Fisher Scientific) in addition to the following cytokines: SCF (300ng/ml), Flt-3L (300ng/ml), IL-3 (60ng/ml) and TPO (100ng/ml), hereafter named “gene therapy media”. All cytokines from Miltenyi Biotech. Flow-sorted cells were cultured in *ex vivo* GT media in a 96 well flat bottom plate coated with 33.3 ug/ml of Retronectin (Takara). The 62 hr protocol consisted of pre-stimulation in GT media (24 hr) followed by transduction with a LV containing GFP (14 hr), an interim incubation in GT media without the vector (10 hr) and a second hit of transduction (14 hr). Vectors of ≥1.19E+08 TU/ml were used for all experiments at multiplicity of infection (MOI) of 100-300. For scRNA-Seq experiments vectors of ≥4.44E+08 TU/ml were used. LV production and purification as well as LV titration are described in **Supplementary Materials**.

### Ethical considerations

the human biological samples were sourced ethically and their research use was in accord with the terms of the informed consents under an IRB/RC approved protocol as specified below.

All animal studies were ethically reviewed by the University of Cambridge Animal Welfare and Ethical Review Body (AWERB) and carried out in accordance with Animals (Scientific Procedures) Act 1986 and the GSK Policy on the Care, Welfare and Treatment of Animals. Animal experiments carried out in Toronto were performed in accordance with institutional guidelines approved by the University Health Network (UHN) Animal Care Committee.

### Primary human samples

All primary human samples were obtained with informed consent from healthy donors by the Cambridge Blood and Stem Cell Biobank (CBSB) in accordance with regulated procedures approved by the relevant research and ethics committees (07/MRE05/44 and 18/EE/0199 research studies). CB samples were pooled independently of sex and processed as a single sample, with the exception of scRNAseq experiments where only single sex CB samples were used. mPB was obtained from healthy male donors aged 25 to 28 by administration of daily Filgrastim (Neupogen) (10 ug/kg per day) for 5 days. Apheresis was performed on day 5 and 6 using the Optia Spectra (Terumo BCT). Experimental work on mPB CD34^+^ cells was performed under 07/MRE05/44 and 18/EE/0199 REC approved research studies.

### Human CB and mPB CD34^+^ cell selection

CB Mononuclear cells (MNCs) were isolated from whole blood (diluted 1:1 in PBS) by Pancoll (PAN-Biotech) density gradient centrifugation at 500g for 25 minutes with the brake off. Red blood cells were lysed by incubation with Red Blood Cell Lysis Buffer containing ammonium chloride for 15 minutes at 4°C (BioLegend). CD34^+^ selection was performed using AutoMACS cell separation technology following incubation with CD34+ selection beads (Miltenyi Biotech) and FcR blocking reagent (Miltenyi Biotech) in PBS + 3% foetal calf serum (FCS) (PAN-Biotech) for 30 minutes at 4°C. CB CD34^+^ cells were then cryopreserved with 20% DMSO (Sigma) in FCS (RBMI) at -150°C until further use. mPB CD34^+^ cells were enriched by the CliniMacs Prodigy system (Miltenyi Biotech) using the automated LP-34 programme for 5 h and 45 minutes consisting of successive washes, CliniMACS® CD34 GMP MicroBeads (Miltenyi Biotec) incubation, immunomagnetic column selection, volume reduction and target cell elution using the TS310 tubing set (Miltenyi Biotec). mPB CD34^+^ cells were cryopreserved in 10% DMSO (Sigma) in FCS (RBMI) at at -150°C until further use.

### Flow cytometry and Fluorescence Activated Cell Sorting

mPB and CB CD34^+^ cells were thawed by dropwise addition of pre-warmed Iscove’s Modified Dulbecco’s Medium (IMDM) (Thermo Fischer Scientific) supplemented with 0.1mg/ml DNAse (Lorne Laboratories) and 50% FCS. Harvested cells were incubated in PBS + 3% FCS containing an antibody mix (**Table S2**; Panel A) for 20 minutes at room temperature (RT). Cells were washed and resuspended in PBS + 3% FCS for cell sorting on the BD FACS Aria Fusion (BD Biosciences) at the NIHR Cambridge BRC Cell Phenotyping hub. Populations were isolated in either a bulk or single cell manner depending on experimental purpose. Single cells were sorted with the single cell purity setting and index data recorded for all surface markers. Bulk cells were sorted with purity setting. Purity for all sorts was estimated at >95%. Previously defined phenotypic populations were sorted: CD19^-^/CD34^+^/CD38-(mPB) and LT-HSC (CD19^-^/CD34^+^/CD38^-^/CD90^+^/CD49f^+^ ; CB and mPB) (*4*) using the representative gates shown in **Fig. S5A**. For scRNA-Seq experiments performed following culture in mPB and CB, phenotypic populations were sorted at 0 h and Zombie- (Live) fractions from each population were re-sorted post-culture at indicated time-points.

Flow cytometry analysis was performed using the BD LSR Fortessa (BD Biosciences), BD LSR Fortessa X-20 (BD Biosciences), FACSCelesta (BD Biosciences), MACSQuant Analyzer 10 (Miltenyi Biotech) and in the case of high throughput analysis the HTS Plate reader was used on the BD LSR Fortessa X-20. Cells were incubated in PBS + 3% FCS containing an antibody mix for 20 minutes at room temperature (RT). All antibodies and antibody panels are listed in **Table S2**.

### Cell cycle assays

For time to first division assays, LT-HSCs were single cell sorted into 96 well u-bottom plates containing 100ul of indicated media per well, centrifuged at 500g for 5 minutes and manually counted every 12 hr for 96 hr using a light inverted microscope. For phoshoRb stainings, cells were harvested at indicated time-points and fixed with neat Cytofix/Cytoperm (BD Biosciences) for 10 minutes at RT. Cells were washed with 400ul of 1x Permwash (BD Biosciences) and stained overnight with Anti-Rb (phospho S807/S811, conjugated to Alexa647 fluorochrome) (Cell Signalling Technologies). Cells were then washed with 1x Permwash and stained with 0.5ug/ml DAPI (ThermoFisher Scientific) in 1x Permwash for 15 minutes. Cells were again washed with 1x Permwash and resuspended in PBS + 3% FCS for flow cytometry analysis. DAPI was recorded on a linear scale and samples analysed at ≤ 30 events/second.

### Cell size measurements

100 CB LT-HSCs per well were sorted into a 384 well plate containing MEM media either untreated (UNTR) or PD treated (200 nM). Cell images were taken every 24 hr in bright field (BF) with a 20x magnification on a Leica DMI300 B microscope using the MetaMorph Microscopy Automation and Image Analysis Software. Cell size was analysed using ImageJ measuring 25 cells/well and expressed as cell diameter (um).

### Mitochondrial activity assay

CB LT-HSCs were cultured in differentiation facilitating conditions, harvested at indicated time-points and stained with the cationic dye 100nM Tetramethylrhodamine (TMRM) (Life Technologies) for 40 minutes (at 37°C & 5% CO2) which accumulates in active mitochondria. Cells were washed and resuspended in an appropriate volume of PBS + 3% FCS for flow cytometry analysis.

### Apoptosis assay

mPB LT-HSCs were cultured in gene therapy conditions and CB LT-HSCs were cultured in differentiation facilitating conditions and harvested at indicated time-points. Harvested cells were washed and resuspended in fridge cold PBS containing Annexin-V/PE (1/20) (BD Biosciences) and 7-AAD (1/20) (BD Biosciences) for 15 minutes at RT. 1X Annexin-V binding buffer (BD Biosciences) was added for immediate flow cytometry analysis.

### Single cell MEM differentiation assay

Cells were single cell sorted into 96 well flat bottom plates containing medium facilitating the growth of myeloid (My), megakaryocyte (Meg), erythroid (Ery) and lymphoid (Lym; only natural killer (NK) cells supported) colonies. This medium comprised of StemPro base media supplemented with Nutrients (0.035%), Pen/Strep (1%), L-Glu (1%), human LDL (50ng/ml) in addition to the following cytokines: SCF (100ng/ml), Flt-3L (20ng/ml), TPO (100ng/ml), EPO (3 units/ml), IL-6 (50ng/ml), IL-3 (10ng/ml), GM-CSF (20ng/ml), IL-11 (50ng/ml), IL-2 (10ng/ml) and IL-7 (40ng/ml). All cytokines from Miltenyi Biotech except EPO (Janssen).Cells were harvested at day 21 of culture into a 96 well u-bottom plate, stained with an antibody mix (**Table S2**, Panel B) for 20 minutes at RT and washed (100ul per well of PBS + 3% FCS) before high-throughput flow cytometry analysis of colony size, colony output and clonogenic efficiency ((number of colonies generated / total single cells plated) * 100). A colony was defined as >30 cells observed in (CD45^+^ & GlyA^+^) gates. Colonies were further assigned to a colony type as follows: Myeloid: > 30 CD45^+^ CD11b^+^ cells; Monocyte: > 30 CD45^+^ CD14^+^ cells; Granulocyte: >30 CD45^+^ CD15^+^ cells; Lymphoid (NK only): > 30 CD45^+^ CD56^+^ CD11b^-^ cells. Undifferentiated colonies were determined as having >30 cells in (CD45^+^ & GlyA^+^ & CD41+) gates but no lineage assignment. Representative gates for analysis are shown in **Fig.S5B**.

### Xenograft transplantation

NOD.Cg-PrkdcscidIl2rgtm1Wjl/SzJ (NSG) mice or NOD.Cg-Prkdcscid Il2rgtm1Wjl Tg(CMV-IL3,CSF2,KITLG)1Eav/ MloySzJ (NSG-SGM3) mice were bred in-house or obtained from Charles River. All experimental cohorts were >11 weeks of age. Only female cohorts were used for experiments involving NSG animals. Both male and female animals were used for experiments involving NSG-SGM3 animals. All animals were housed in a specific pathogen free animal facility and experiments were conducted under UK Home Office regulations or in accordance with institutional guidelines approved by the UHN Animal care.

For primary transplantation experiments NSG mice were sub-lethally irradiated (2.4 Gy) 24 h prior to transplantation. For intrafemoral (IF) injections, mice were anesthetised with isoflurane and transplanted with the indicated cell dose in PBS + 0.1% Pen/Strep (ThermoFisher) (25ul). Following transplantation, mice were injected subcutaneously with the analgesic buprenorphine (Animalcare) at 0.1mg/kg. For intravenous injection, mice were transplanted with a cell suspension (max 150ul volume) in PBS + 0.1% Pen/Strep by tail vein injection. For all xenograft experiments involving cell culture, injected cell doses at time-points are representative of the cell count at the time of sort (0 h). Mice were culled and bone marrow harvested 18-20 weeks for primary transplantation experiments. The femur and tibia bones from the two hind legs were taken and for IF injected mice, the injected femur was analysed separately. Bone marrow was stained in an antibody panel (**Table S2**; Panel C) for 20 minutes before washing and resuspension in PBS + 3% FCS for flow cytometry analysis. Methods related to secondary transplantation are reported in **Supplemental Materials**.

For all studies, mice were considered engrafted if human cells (hCD45^++^ &GlyA^+^) >0.01% of Singlets and if >20 cells were present in any lineage determination gate (**Fig.S5C**).

### scRNA-Seq library preparation

Single cell RNA-Sequencing (scRNA-Seq) libraries were prepared using a published Smart-Seq2 protocol (*37*), adapted as described in **Supplemental Materials**. Libraries were quantified using KAPA library quantification kit (Roche) and 20nM of libraries sequenced by paired end sequencing (150bp) using Illumina HiSeq4000 (Illumina) and NovaSeq 6000 (Illumina) at the CRUK-CI genomics core facility (Cambridge, UK).

### scRNA-Seq experimental design

Two main scRNA-Seq datasets were generated in this study: a time-course of LT-HSCs cultured in EXPER conditions (Dataset 1), a time-course of LT-HSC cultured in GT conditions (Dataset 2). Dataset 1 was acquired over 2 independent experiments (Batch 1 and 2) using LT-HSCs isolated from CB samples from 2 independent pools of male donors at the timepoints and conditions indicated in **Table S3**. Dataset 2 was acquired over 4 independent experiments (Batches 1 to 4) using LT-HSCs isolated from mPB samples from 4 independent healthy male donors at the timepoints and conditions indicated in **Table S4**. UNTR and PD (200nM) treated conditions were always paired in the same batches.

Two main integrations were performed in this study: i) all CB LT-HSCs from 0h and untreated timepoints of Dataset 1 (Integration 1); ii) all single cells of Dataset 1 and Dataset 2 (UNTR and PD treated; Integration 2). Integration 1 was performed with 2 independent methods: i) Seurat 4 method (described below); ii) ScanPy method (described in **Supplementary Materials**). Integration 2 was performed with the Seurat 4 method.

### scRNA-Seq quality control

Read alignment was performed using GSNAP (*77*) against Ensembl genes and initial quality control (QC) was performed by FastQC (*78*). The count matrix was generated by HTSeq (*79*). Additional QC was then performed in the bglab package (*80*) using the determined thresholds shown in **Table S5**) yielding the number of cells passing QC reported in **Table S3** for Dataset 1 and **Table S4** for Dataset 2.

### Seurat 4 pipeline for batch correction and batch/dataset integration

Seurat objects were created using the Seurat package (v4) (*81*). Genes from raw counts were filtered if not detected in > 3 cells. The function SCTransform was used to perform normalisation and variance stabilisation, the mitochondrial genes percentage was regressed out. All other parameters were left as default if not mentioned specifically. For cell cycle regression, S and G_2_-M cell cycle scores were calculated using the CellCycleScoring function on the object following SCTranform. The values of G_2_-M scores were subtracted from the values of S scores resulting in the difference of the cell cycle scores. SCTransform was applied on the Seurat object again regressing the mitochondrial genes percentage and the difference of the cell cycle scores. For batch/dataset integration, Seurat objects to be integrated or batch corrected were curated into a list. 3,000 features were selected with the function SelectIntegrationFeatures. The list of objects was prepared to integrate using the function PrepSCTIntegration. FindIntegrationAnchors function was used to find a set of anchors for integration, the k.filter parameter was set to 100. Integration was performed with the function IntegrateData. PCA was computed on the batch corrected/aligned Seurat objects with the RunPCA function. UMAPs were computed using the RunUAMP function with the dims parameter set to 1:30.

### Pseudotime analyses

The Monocle3 package (version 1.2.9) (*82*) was used to find the pseudotime ordering of the samples. Seurat integrated objects were converted to Monocle3 objects with the function as.cell_data_set from the package seurat-wrappers (*83*). The cluster_cells function was run with the parameter reduction_method set to ‘UMAP’. Principal graph was learned with the function learn_graph (with use_partitition parameter set to TRUE for Integration 1 and set to FALSE for Integration 2). Cells were ordered on the principal graph using the order_cells function, with a manually chosen root cell.

### Differential expression and definition of “ex vivo modulated genes”

Genes from the raw count matrix were filtered if expressed in < 4 cells. Filtered raw counts were used to perform differential expression analysis with the R package DESeq2 (version 1.36.0) (*84*). Batch effects were accounted for in the model where applicable. For Integration 1, the union of all differentially expressed genes each pairwise time-point comparison was curated, generating a list of 10,010 genes, herein referred to as “ex vivo modulated genes” (**Data S4**). A variance stabilizing transformation was performed in the DESeq2 package from the differential expression analysis. Batch effects were removed on the variance stabilised matrix using the limma package correction (version 3.52.2) (*85*). These VST batch corrected values were used for visualisations of gene expression in violin plots and as input for DegPatterns.

### Gene set enrichment and variation analysis

Gene Set Enrichment Analysis (GSEA) (v4.2) (*86*) was performed against the c2 curated pathway database using a pre-ranked list by the stat value (value of the Wald test statistic) from the DESeq2 output specifying 10,000 permutations. Gene-Set Variation Analysis (GSVA) was also performed (*40*) against c2 curated pathways using the normalised, batch corrected count matrix generated by Scanpy (parameters min.siz=10, max.sx=500). GSEA and GSVA were also performed with curated signatures. All gene signatures were created using the top 100 differentially expressed genes contrasting previously reported populations. Signatures were curated from CD34^lo^CLEC9A^hi^ (Subset1) and CD34^hi^CLEC9A^lo^ (Subset 2) from (*19*) and from human dormant and activated HSC populations (*22*).

Other experimental methods (LV production and purification, LV vector titration, adaptation of the Smart-Seq2 protocol, secondary transplantation in xenografts) as well other bioinformatic methods (Scanpy normalisation and processing of counts, identification of gene expression along the time course, scEntropy measurements, Bayesian modelling of gene expression and over-dispersions to measure expression variability, correlations of median expression) are reported in **Supplementary Materials**.

### Statistical analysis

Analysis of the HSC frequency (%LTRC) from transplanted populations was performed by using Extreme Limiting Dilution Analysis (ELDA) software (https://bioinf.wehi.edu.au/software/elda) taking into account the number of engrafted mice, the total mice used and the cell dose transplanted. For analysis of a statistical difference in the HSC frequency within two groups, a Chi-Squared test was performed within the ELDA software. Flow cytometry data was analysed using FlowJo software (v10). For analysis of colony data derived from single sorted LT-HSCs, FlowJo v.10 gating statistics were exported and data further analysed in R Studio (v.1.2). Graphpad Prism (v9.3), python (v3.8.6) and R Studio (v1.2) were used for the creation of all plots. Statistical analysis between multiple groups was performed in Graphpad Prism or R Studio. Normality of data was deduced from the Shapiro-Wilks normality test. For statistical analysis between two groups a parametric test (Students t-test) or a non-parametric test (Mann-Whitney U-test) was performed. For analysis between multiple groups, an analysis of variance (ANOVA) test was performed. All statistical tests were performed with a confidence interval of 95%.

## Acknowledgments

We would like to thank the CB and mPB donors for their kind donation of human tissue, the Cambridge Blood and Stem Cell Biobank, specifically Dr Joanna Baxter and the nurses who consented and collected CB samples; the Cambridge NIHR BRC Cell Phenotyping Hub for their flow cytometry services and advice, and the CRUK Cambridge Institute genomics centre for sequencing.

## Funding

Wellcome – Royal Society Sir Henry Dale Fellowship 107630/Z/15/Z (EL)

European Hematology Association Non Clinical Research Fellowship Award RG20 (EL)

Core support grants by Wellcome and Medical Research Council (MRC) to the Wellcome-MRC

Cambridge Stem Cell Institute 203151/Z/16/Z (EL, BG, ARG)

GSK-Varsity Alliance (NF, EL, BG)

MRC iCASE PhD studentship 1942750 (CSJ)

Cancer Research UK Cambridge Cancer Centre PhD fellowship (SB)

Deutsche Forschungsgemeinschaft Research Fellowship ME 5209/1-1 (NM)

Cancer Research UK C1163/A21762 (BG, ARG)

Blood cancer UK 18002 (BG)

Gates Cambridge Scholarship, Gates Cambridge Trust (MD)

Alborada Trust (ARG)

William B Harrison Foundation (ARG)

University of Toronto’s Medicine by Design initiative with funding from the Canada First Research Excellence Fund (JED)

## Author contributions

Conceptualization: EL, NF, CSJ, SB, BG

Methodology: CSJ, SB, WL, NKW

Investigation: CSJ, SB, WL, KBK, GK, EFC, JML, SS, CG, MD, NF, MJW

Visualization: KS, XW, CSJ, SB, EL

Formal analysis: KS, XW, ED

Data curation: KS, XW

Validation: EL, KS, CSJ

Resources: NF, SH, CB

Writing—original draft: CSJ, KS, EL

Writing—review & editing: CSJ, KS, EL

Supervision: EL, NF, BG, JED, CB, NKW, ARG

Project management: EL, NF

Funding acquisition: EL, NF

## Competing interests

The authors declare that they have no competing interests.

## Data and materials availability

Sequencing files and metadata associated to Smart-Seq2 datasets are deposited at GEO with accession numbers GSE213365 (time course of CB LT-HSC cultured in differentiation medium) and GSE213370 (time course of mPB cultured in GT medium) under the super series GSE213372. All code is publicly available on GitHub at https://github.com/elisa-laurenti/LT-HSC_exvivo.

All data needed to evaluate the conclusions in the paper are available in the main text or the Supplementary Materials. All other data or materials will be distributed by the authors upon request.

## Notes

https://github.com/elisa-laurenti/LT-HSC_exvivo

### Competing Interest Statement

The authors have declared no competing interest.

### Summary of Updates

Figure 1H and Table S1 revised to updated analysis.

