## Supplementary Material for "Adaptation to *ex vivo* culture drives human haematopoietic stem cell loss of repopulation capacity in a cell cycle independent manner"

**This PDF file includes:**

Supplementary Text (extended description of experimental and bioinformatic methods)
Figs. S1 to S5
Tables S1 to S5
References (87 to 90)
Legends to Data S1 to S7

### Supplementary Text

Extended description of experimental methods

**Lentiviral vector (LV) production and purification.** Third generation self-inactivating (SIN) lentiviral vectors expressing GFP under control of an EIF1$\alpha$ promoter were produced by transient transfection in HEK293T cells. HEK293T cells were cultured at 37 °C/ 5% CO2 in an orbital shaker and passaged until an optimum density ($1.5x10^6$ cells/ml) achieved. Harvested HEK293T cells were transfected with four plasmids (incorporating Gag-Pol (pG3-SYNGP), Rev (pG3-REV), VSVG (pG3-VSVG) and the transfer plasmid containing an eGFP construct (pG3T)) in GMP grade Stem Cell Growth Medium (SCGM) (CellGenix) with Polyethylenimine PEIPro (Polyplus) added to enhance transfection. Plasmids were obtained from the GSK research facility (Stevenage, UK). Cells were incubated (37 °C & 5% CO2) and 5mM of Sodium Butyrate (Sigma) was added to enhance transfection at 24 h. HEK293T cells were harvested at 72 h post transfection and LV containing medium was separated from cell debris by centrifugation at 1000g for 20 minutes at 4 °C with the supernatant clarified using 0.8um and 0.45um vacuum filters (Corning). LV particles were concentrated by ultracentrifugation at 5000g for 20 h at 4 °C the supernatant was discarded and the pellet air dried. Pellets were resuspended in SCGM and stored at -80°C.

**LV titration.** CEM A3-04 cells at optimum density ($1.5x10^6$ cells/ml) were resuspended in RPMI (Thermo Fisher) containing 8µg/ml protamine (Sigma) and plated at a density of $1.5x10^6$ cells/ml (density kept constant between wells). Serial dilutions (4-1 to 4-10) of purified LV were added to wells and cells incubated (37 °C & 5% CO2) for 2 h. Additional RPMI 1640 supplemented with 1% L-Glu, 10% FCS and 1% Penicillin/Streptomycin (Pen/Strep) (ThermoFisher Scientific) was added to each well and plates incubated (37 °C & 5% CO2) for 4 days. Harvested CEM cells were resuspended in PBS + 5% Human Serum Albumin (HSA) (Irvine Scientific) containing 1% 7-AAD (Biolegend). GFP positivity was analysed on the MACSQuant Analyzer 10 (Miltenyi Biotech). Vector titre (TU/ml) was calculated as = (Number of cells x %GFP+ x Dilution Factor) / Transduction volume (0.5 mL).

**Smart-Seq2 adapted protocol.** Single cell RNA-Sequencing (scRNA-Seq) libraries were prepared using an adapted Smart-Seq2 protocol (*37*). A lysis buffer was prepared of 20U/ul SUPER-In RNase inhibitor (Thermo Fisher Scientific) and 0.4% Triton-X100 (ratio of 1:19). The lysis buffer was added to a mix of DTT (Thermo Fisher Scientific) (final concentration of 0.57mM), dNTPs (Invitrogen) (final concentration 0.11mM) and nuclease free water (Thermo Fisher Scientific) and 4ul aliquoted per well in a 96 well PCR plate and stored at -80°C. Upon thawing and single cell sorting, an annealing mix was added (containing final concentration of External RNA Consortium Controls (ERCC) RNA Spike-In mix diluted $1:3x10^6$ (Thermo Fisher Scientific), 10uM Oligo-dT30 VN (Thermo Fisher Scientific) and nuclease free water (ThermoFisher Scientific) and annealing performed (3 minutes). The plate was briefly centrifuged (1000g for 30 seconds at 8°C) to collect liquid at the bottom of the well, the reverse transcription master mix added (containing 10U/µl SMARTScribe reverse transcriptase (Thermo Fisher Scientific), 1U/ul SUPER-In RNase inhibitor (Thermo Fisher Scientific), 2uM Template Switching Oligo (TSO), 5X First Strand Buffer (Thermo Fisher Scientific) and nuclease free water (Thermo Fisher Scientific)) and reverse transcription performed (~2.5 h). The plate was again briefly centrifuged (1000g for 30 seconds at 8°C), the PCR master-mix added (containing 166nM

IS-PCR primer, 2X KAPA HiFi HotStart ReadyMix (Roche) and Nuclease free water) and 23 cycles of PCR performed to account for the low RNA content of quiescent HSCs (~3 h). Plates were stored at -20°C until PCR purification. RT Ampure XP Beads (Beckmann Coulter) were mixed and incubated with PCR product (1:0.6/0.7 ratio) and PCR purification performed as in (*37*), to remove fragments of a small, suboptimal length. The Biomek FXP Automated Workstation was used for experiments involving >3 plates. The quality of material was checked using the Agilent High Sensitivity DNA kit (Agilent) following manufacturer's instructions. The Quant-iT PicoGreen dsDNA kit and reagents (Thermo Fisher Scientific) were then used to determine the concentration of DNA (triplicate values per plate per cell condition) and dilution plates created (using Elution buffer as diluent) for all samples to achieve 0.1-0.15ng/ul. Tagmentation was performed (10 minutes) using the Nextera XT DNA library preparation kit (Illumina) mixing 2.5ul Tagment DNA Buffer, 1.25ul Amplicon Tagment Mix and 1.25 ul of DNA per sample. The Tn5 transposase was stripped using NT buffer (Illumina) (1.24ul per sample) and a PCR mix added (containing 3.74ul of NPM mix (Illumina) 1.24ul of appropriate i7 and 1.24ul of i5 primers (Illumina) per sample) with PCR performed (~30 mins) to amplify adapter-ligated sequences. PCR product purification was performed in two successive steps using RT Ampure XP beads (1:0.5, 1:0.3) to yield final average amplicon lengths of ~400-700 bp.

**Secondary transplantiation in xenografts.** For secondary transplantation experiments in the EXPER system, NSG-SGM3 mice were irradiated using 2.25Gy 24 hr prior to transplantation and primary BM samples were thawed in X-VIVO 10 media (Lonza) + 50% fetal bovine serum (Wisent) supplemented with DNase (100 µg ml−1, Roche). Viable (SytoxBlue-) (ThermoFisher Scientific) human CD45++ cells were sorted (**Table S2**; Panel D) on the Aria Fusion (BD Biosciences). Cells were pooled based on condition and intrafemorally injected in three doses. Mice were culled at 8 weeks, bone marrow harvested and stained in an antibody panel (**Table S2**; Panel E) and analysed by flow cytometry with the same methodology as in primary transplantation experiments. Secondary transplantation experiments for EXPER conditions were kindly performed at UHN, Toronto.

For secondary transplantations of experiments in the GT system, primary mouse BM was thawed by dropwise addition of pre-warmed Iscove's Modified Dulbecco's Medium (IMDM) (Thermo Fisher Scientific) supplemented with 0.1mg/ml DNAse (Lorne Laboratories) + 50% FCS. Cells were counted and injected in three doses by IV injection (max 150ul volume) in PBS + 0.1% Pen/Strep. Mice were culled at 8 weeks, bone marrow harvested and stained in an antibody panel (**Table S2**; Panel C) and analysed by flow cytometry with the same methodology as in primary transplantation experiments.

### Extended description of bioinformatic methods

**Scanpy normalisation and processing of counts.** Raw counts of both batches from Dataset 1 were processed using the Scanpy package (version 1.4.5.1) (*87*). Scanpy anndata objects of both batches were created and concatenated. The function filter_genes was run with parameter min_cells=3. After filtering, 33,774 genes were left. Cells were normalised with the function normalize_total with parameter target_sum=1e4. Counts were log transformed with a pseudocount of 1 added to mitigate the mean-variance relationship using the log1p function, to reduce skewing of data and account for drop-outs. Batch effects were regressed out with the function combat, with

the incubation time of the samples set to be the covariates. The scanpy combat function is a wrapper function for the combat package (https://github.com/brentp/combat.py).
Highly variable genes were selected with the function highly_variable_genes, which implements the works of (*88*). The parameters for the highly_variable_genes function were as follows: min_mean=0.05, max_mean=13, min_disp=0.1, max_disp=3. The anndata object was subset with only 9,212 highly variable genes. Principal component analysis was performed using the pca function in the Scanpy package with default parameters. Nearest neighbours for the cells were selected using the bbknn package (version 1.3.7) (*89*). The function bbknn_pca_matrix was used with the PCA matrix calculated by Scanpy and the batch information as inputs, parameters were set as follow: approx=False, metric='euclidean'. The function umap was used to find the UMAP representation of the data, the parameter n_components=3.

**Identification of patterns of gene expression along the time course.** The degPatterns function from the DEGreport package (*39*) was used to group genes based on expression pattern, inputting VST limma corrected matrix values for *ex vivo* modulated genes. The time parameter was set to the incubation time of the samples and eachStep=TRUE. 8,966 genes remained after degpatterns filtering of clusters with <15 genes. A distance matrix was generated from all pairwise comparisons between time-points and hierarchical clustering was performed specifying 13 clusters. Upon visual observation 2 of the 13 clusters were manually split generating 15 clusters which fit the dataset without showing redundancy in expression pattern (clusters were renamed 1-15 accordingly).

To group pathway patterns based on expression trend, the degPatterns function from the DEGreport package (*39*) was again used, inputting the GSVA score matrix, with the parameter time set to the incubation time of the samples and eachStep=TRUE. 4,367 out of 4,596 pathways remained after filtering and 13 clusters again were specified to fit the dataset. Upon visual observation, 6 of 13 clusters of the GSVA score patterns were manually split further, resulting in 19 clusters with no redundancy in expression pattern observed (clusters were renamed 1-19 accordingly).

**scEntropy measurements.** To measure the transcriptomic order of single cells in the dataset, the package scEntropy was used (*90*). The scEntropy per cell is defined as the difference between the cell of interest and an intrinsic reference value calculated by the package (parameter option =RCSA). Scanpy was used to pre-process raw counts from both batches separately and cells were normalised by $1x10^4$ reads (normalize_total function with parameter target_sum=1e4). The scEntropy value for single cells was calculated. To account for batch effects, the difference in the mean entropy values of batch 1 and batch 2 was substracted from each entropy value of batch 1.

**Bayesian modelling of gene expression and over-dispersions to measure expression variability.** The BASiCS package (version 2.2.4) (*46*) was used to perform Bayesian modelling of the counts. The amount of ERCC molecules present together with each cell was calculated from the concentration of the ERCC mix added and the information for the ERCC mix acquired online ([https://assets.thermofisher.com/TFS-Assets/LSG/manuals/cms_095046.txt]). Raw counts were filtered to average reads per million is > 20, leaving 8,464 genes. BASiCS objects were created with the filtered counts, ERCC information and batch information. The objects were created independently per incubation time sample. For the 24h time sample, one cell was not included for

the creation of BASiCS object due to low quantity of ERCC present. Bayesian inference of the parameters of the distributions of the gene counts was performed using the function BASiCS_MCMC. The parameters for BASiCS_MCMC function which calculate residual over dispersion (ε)) were as follow: N=10000, Thin=10, Burn=1000, WithSpikes=TRUE and Regression=TRUE.
To identify residual over-dispersed genes for all incubation time samples, the function BASiCS_TestDE was used that compares the value of residual over-dispersion and performs statistical testing when assessing values between 2 samples. Genes that are differentially over-dispersed based on the residual over-dispersion value were curated for each incubation time sample. To calculate maximally variable genes at each incubation time, genes that were differentially variable in multiple pairwise comparison were assigned to the incubation time with the highest residual over-dispersion value.

**Correlations of median expression.** For EXPER conditions, the median expression value for *ex vivo* modulated genes plus genes differentially expressed in PD treated comparisons (genes changed in 0 h vs 72 h PD, 72 h PD vs 72 h UNTR, 0 vs 24 h PD, 24 h PD vs 24 h UNTR) (n=10,903 genes total) was plotted for each comparison and the Pearsons correlation co-efficient calculated (95% CI calculated; mean value shown). For GT conditions, the median expression value for the sum of genes changed in 0 h vs 62 h UNTR, 0 h vs 62 h PD and 62 h PD vs 62 h UNTR, was used (5,469 genes) and each cell at each time-point was compared. All gene lists can be found in **Data S4.**

**Visualisation:** data graphs were produced in R, Python or GraphPad. Figures 4C and S3B were created with BioRender with appropriate licensing options.

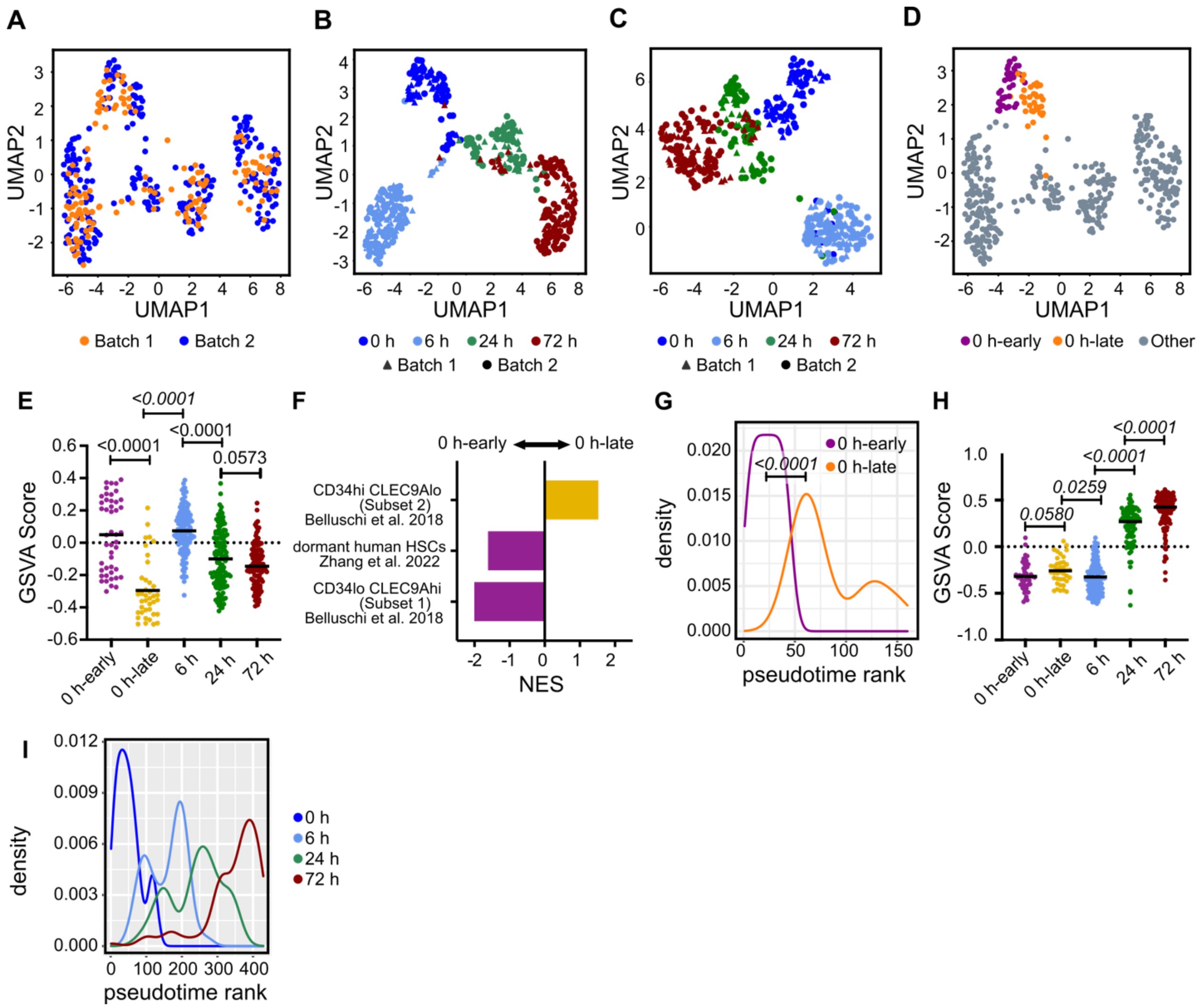


**Fig. S1. Additional bioinformatic analyses of time course scRNAseq data of cultured LT-HSCs and characterization of heterogeneity of quiescent LT-HSCs**

**(A-D)** UMAPs of 429 single LT-HSCs cultured in the EXPER system over a time-course of 0 h, 6 h, 24 h and 72 h. **(A)** UMAP coloured by experimental batch and generated using Seurat 4 pipeline following cell cycle regression. **(B)** UMAP coloured by time-point with shape representing experimental batch and generated using Seurat 4 pipeline with no cell cycle regression. **(C)** UMAP coloured by time-point with shape representing experimental batch and generated using Scanpy pipeline with no cell cycle regression. **(D)** UMAP as in (A) showing k=2 clusters defined within 0 h cells: 0 h-early (purple, n=46 cells) and 0 h-late (yellow, n=39 cells).

**(E)** GSVA scores per cell of “KRIEG_HYPOXIA_VIA_KDM3A” geneset in the indicated conditions. Unpaired t-test.

**(F)** GSEA analysis comparing 0 h-early and 0 h-late LT-HSCs to curated signatures generated from dormant human HSCs (*21*), multipotent LT-HSCs (CD34hi CLEC9Alo; Subset 1) and myelo-lymphoid restricted LT-HSCs (CD34$^{lo}$ CLEC9A$^{hi}$; Subset 2) (*18*), all FDR < 0.05.

**(G)** 2D Pseudotime rank plot of 0 h-early and 0 h-late. Wilcoxon rank sum test performed.

**(H)** GSVA scores per cell of "REACTOME_RESPIRATORY_ELECTRON_TRANSPORT_ATP_SYNTHESIS_BY_CHEMIOSMOTIC_COUPLING_AND_HEAT_PRODUCTION_BY_UNCOUPLING_PROTEINS" geneset in the indicated conditions. Unpaired t-test.

**(I)** 2D Pseudotime rank plot of LT-HSCs over time-course generated with no cell cycle regression applied.

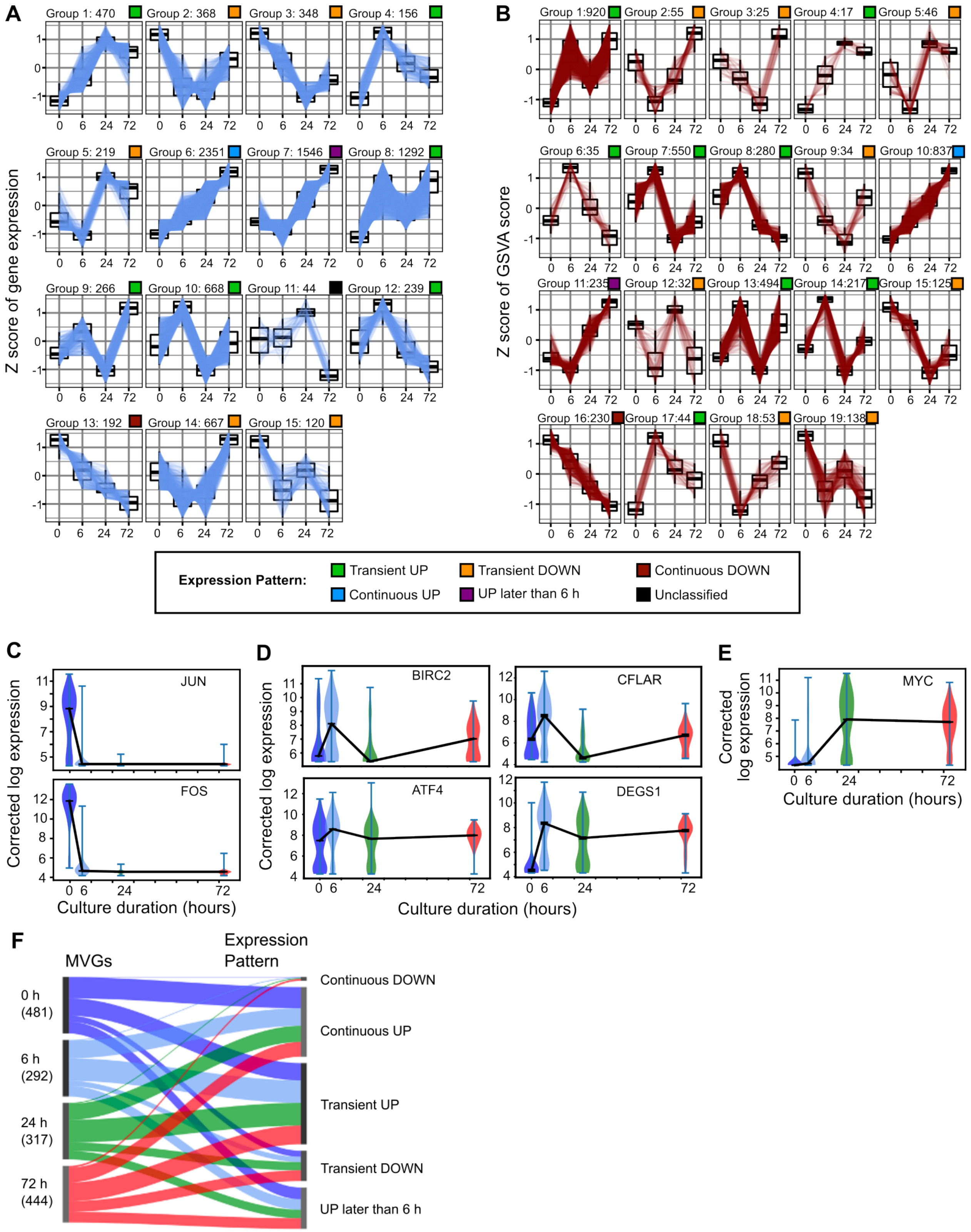


**Fig. S2. Dynamic patterns of LT-HSC gene expression observed during *ex vivo* culture.**

**(A-B)** Output from DEG Report clustering of gene expression patterns **(A)** or GSVA pathway patterns **(B).** After manual curation of the respective DEG Report outputs, 15 gene expression patterns **(A)** and 19 GSVA pathways patterns **(B)** were retained, which were found to fit the dataset without showing redundancy in expression pattern. **(A)** 10,010 genes classified total; 8,966 genes after filtering. **(B)** 4,596 total GSVA pathways classified; 4,367 pathways after filtering. **Data S5**.

**(C-E)** Violin plot of corrected log expression values for selected genes. AP-1 transcription factors *JUN* and *FOS* **(C)**, apoptosis master regulators *BIRC2* and *CFLAR* **(D)**, regulators of proteostatic stress *ATF4* and *DEGS1* **(D)**, and transcription factor *MYC* **(E)**. Line connecting median values between time-points.

**(F)** Sankey plot of maximally variable genes (MVG) at each time-point (left) matched to corresponding expression pattern (right) (total MVG at 0 h = 853; 6 h = 470; 24 h = 605; 72 h = 864 genes).

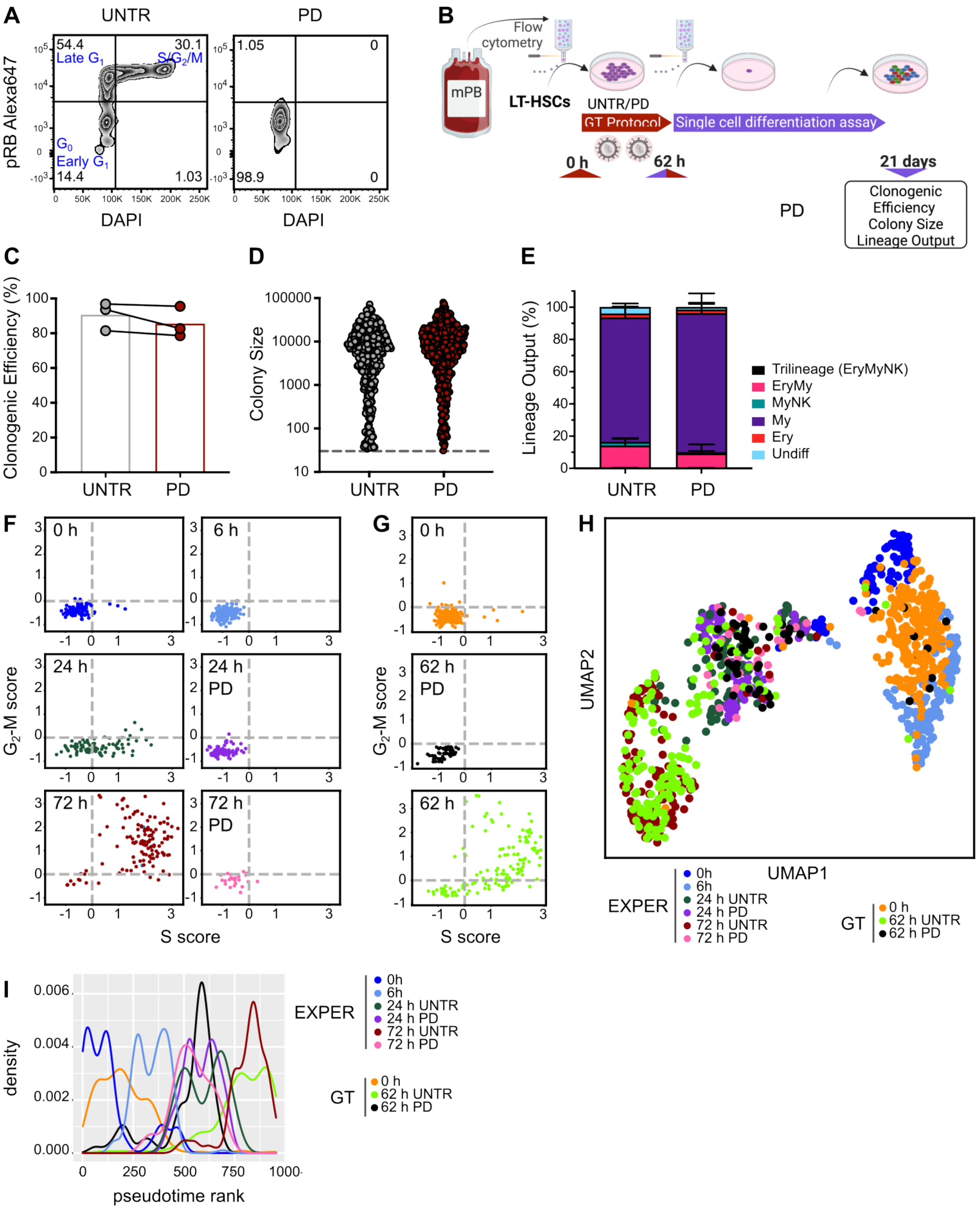


**Fig. S3. Validation of reversible block of LT-HSCs in early $G_1$ by PD and its transcriptional effects**

**(A)** Representative example of pRb/DAPI flow cytometry plot of alive UNTR (left) or PD treated (right) mPB LT-HSCs cultured for 62 h in GT conditions.

**(B)** Workflow of GT protocol culture and single cell differentiation assay. mPB LT-HSCs were flow cytometry sorted in bulk and cultured in GT protocol conditions for 62 h including two rounds of LV transduction. Live (Zombie$^-$) cultured LT-HSCs were then single cell sorted and cultured for an additional 21 days in a single cell differentiation assay in media supporting the generation of Myeloid, Erythroid and Megakaryocytic colonies (MEM). At Day 21 colonies were analysed for clonogenic efficiency, colony size and lineage output. Created with BioRender.

**(C-E)** Clonogenic efficiency ((number of wells generating true colonies / number of wells plated)*100) **(C)**, colony size distribution **(D)** and types of colonies (lineage output, Ery = CD45$^-$ GlyA$^+$; My = CD45$^+$CD14$^+$; NK = CD45$^+$CD56$^+$CD11b$^-$)) **(E)** of mPB LT-HSCs UNTR/PD treated cultured in GT protocol followed by culture in single cell MEM differentiation assay. A colony is defined by >30 cells in CD45$^+$ & GlyA$^+$ gates (n=3 biological repeats, UNTR = 421 colonies and PD treated = 369 colonies).

**(F-G)** Transcriptional allocation of cell cycle status of UNTR/PD LT-HSCs cultured in EXPER medium (**F**, n=2 experimental batches; n=536 total cells) or in the GT system (**G**, n=4 experimental batches; n=418 total cells).

**(H)** UMAP visualisation of scRNAseq from 954 LT-HSCs from the indicated culture conditions (EXPER: 536 single cells: GT: 418 single cells). No cell cycle regression applied.

**(I)** 2D Pseudotime density rank plot of LT-HSCs cultured in EXPER and GT systems. No cell cycle regression applied. Number of cells as in (H).

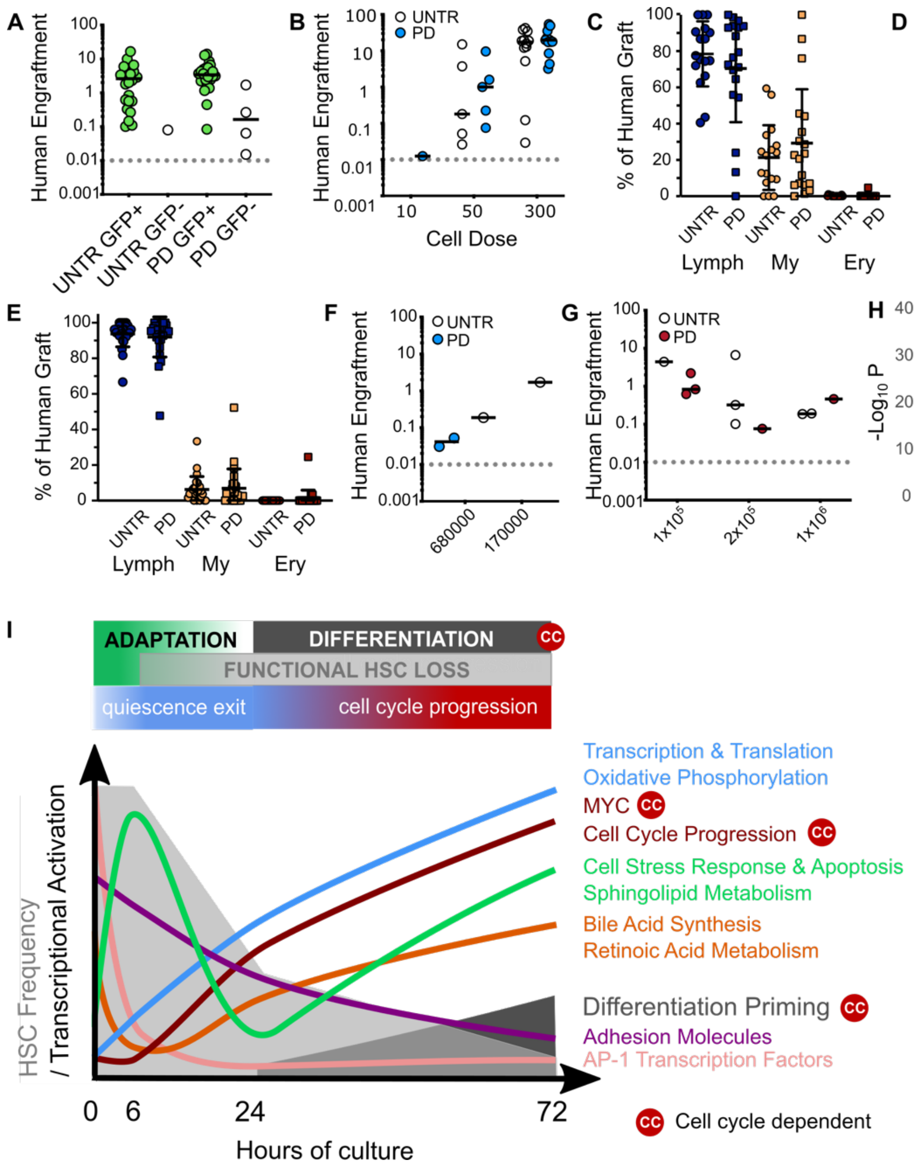


**Fig. S4. PD treatment *ex vivo* does not affect engrafting capacity of LT-HSCs but alters the expression of genes linked to differentiation**

**(A)** Graft size in mice engrafted with mPB $CD34^+/CD38^-$ split by GFP status ($GFP^+$ graft defined as ≥30 cells in $GFP^+$ gate). $GFP^+$ grafts were observed in 24/25 mice engrafted with UNTR cells (96%) and 22/26 of mice engrafted with PD treated cells (85%).

**(B-C)** Graft size (% of human $CD45^{++}$ and $GlyA^+$ ) **(B)** and lineage output **(C)** at 18 week post transplantation of UNTR/PD treated LT-HSCs cultured for 24 h in EXPER system (n=3 biological experiments; n= 31 UNTR mice and n=30 PD treated mice). Graft size in 300 cell dose of PD compared to UNTR: unpaired t-test *p=0.3423*. Lineage output: UNTR Myeloid compared to PD treated Myeloid: Mann Whitney U test: *p=0.5577*). Lymphoid = $CD45^{++}CD19^+$ ; Myeloid = $CD45^{++}CD33^+$ ; Erythroid = $CD45^-GlyA^+$.

**(D)** Lineage output after long-term (18 week) transplantation of UNTR/PD treated LT-HSCs cultured for 72 h in EXPER system (n=5 biological experiments; n=42 PD mice, n = 38 UNTR mice).

**(E)** Lineage output at 18 week post-transplantation of UNTR/PD treated mPB $CD34^+/CD38^-$ cultured for 62 h in GT system (Mann Whitney U-test UNTR Erythroid vs PD treated Erythroid $p = 0.4902$).

**(A-E)** Raw data available in **Data S1**.

**(F-G)** Graft size in secondary transplantation experiments from primary recipients initially engrafted with UNTR/PD treated LT-HSCs cultured for 72 h in EXPER system **(F)** (n= 20 mice total; 10 UNTR, 10 PD; n=1 experiment) and UNTR/PD treated mPB $CD34^+/CD38^-$ cells cultured for 62 h in GT system **(G)** (n=21 mice total; UNTR = 11 mice; PD = 10 mice; n=1 experiment)

**(H)** Violin plot of GMP signature genes differentially expressed (FDR<0.05) between CB LT-HSCs cultured for 72h in presence or absence of PD (17 genes shown).

**(I)** Graphical summary of the findings of this study. CC demarks processes that are dependent on cell cycle progression.

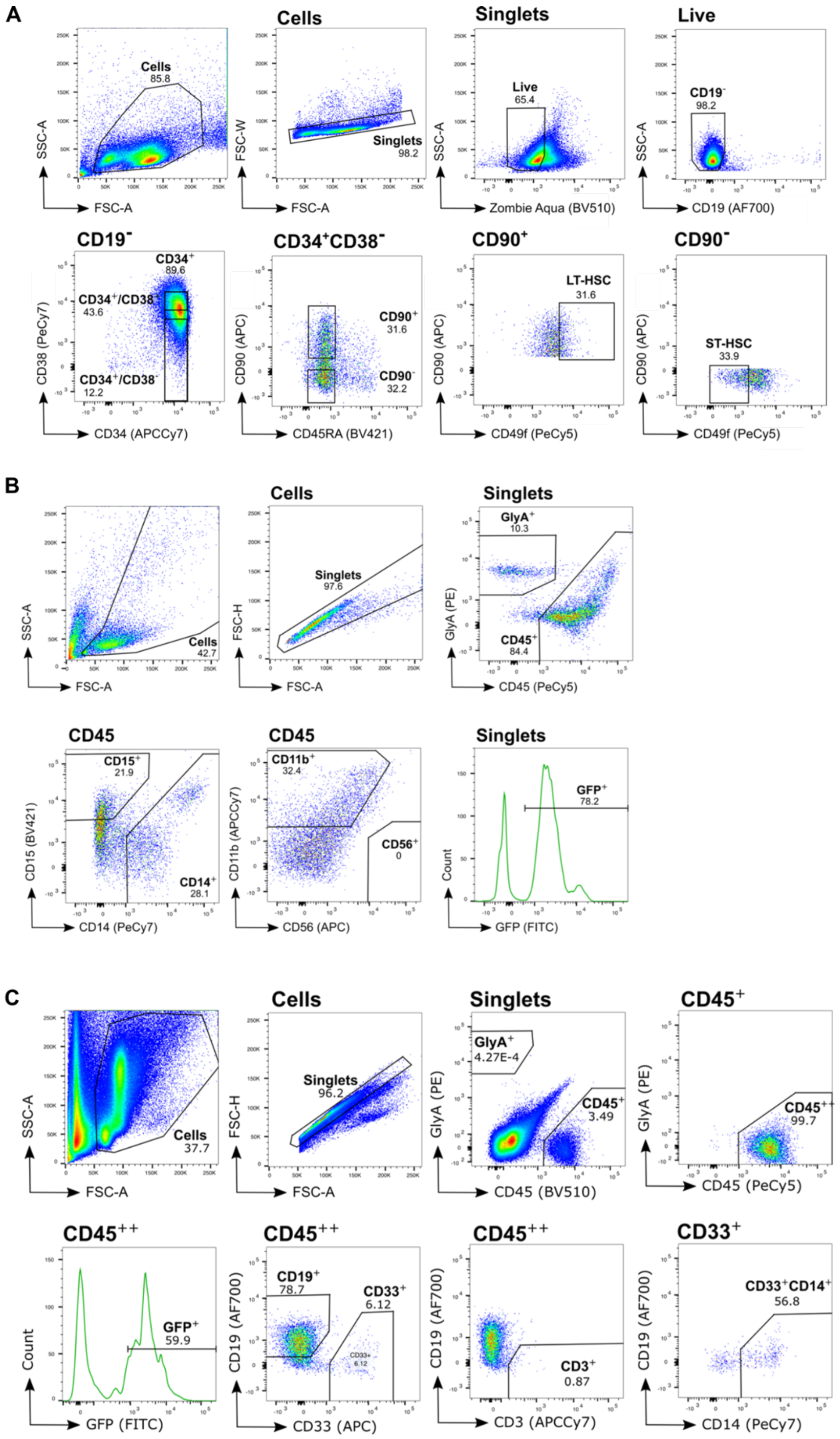


**Fig. S5. Representative gates for flow cytometry experiments**

**(A)** Representative gating strategy for isolation of LT-HSC. The example is here is from a 0 h mPB $CD34^+$ enriched donor sample. LT-HSC: $Zombie^-/CD19^-/CD34^+/CD38^-/CD90^+/CD49f^+$ (top 30% of CD90/CD49f expression). The same strategy was used for CB LT-HSCs.

**(B)** Representative gating strategy for determination of colony size and lineage output from single cell MEM assay. Representative gating of a colony arising from a single sorted mPB LT-HSC after 21 days culture in MEM media. A true colony was determined if >30 cells are observed in each of the $CD45^+$ and $GlyA^+$ gates. Colonies were assigned to specific lineages if >30 cells were found in the following gates: myeloid: $CD45^+/CD11b^+$; monocyte: $CD45^+/CD14^+$; granulocyte: $CD45^+/CD15^+$; lymphoid (NK only): $CD45^+/CD11b^-/CD56^+$. Undifferentiated colonies were determined as having >30 cells in ($CD45^+$ & $GlyA^+$) gates but <30 cells in all the other lineages.

**(C)** Representative gating strategy for analysis of mouse BM after primary transplantation (18-20 weeks). The example here is from a mouse injected with mPB $CD34^+/CD38^-$ cells cultured in GT conditions (62 h) and transduced with a LV containing GFP. Erythroid (Ery) cells were identified as $CD45^-/GlyA^+$. Myeloid (My) cells were identified as $CD45^{++}/CD33^+$. Lymphoid (Lym) cells were identified as $CD45^{++}/CD19^+$. Mice were considered engrafted if ($hCD45^{++}$ &$GlyA^+$) >0.01% of singlets and a lineage detected by containing >20 cells in any respective lineage gate. Same gating strategy was used for CB experiments and all secondary transplantation experiments.

| Culture conditions | Cell type | Hours of culture (h) | Cell Dose | Number of tested mice | Number of engrafted mice | Number of experiments | LTRC frequency (ELDA estimate and 95% CI ) | % LTRC (ELDA estimate and 95% CI ) |
|---|---|---|---|---|---|---|---|---|
| **GT** | $CD34^+$ $CD38^-$ | 0 | 8000 | 3 | 3 | 2 | 1 in 838 (415-1691) | 0.119 (0.059-0.241) |
| | | | 5000 | 5 | 4 | | | |
| | | | 1250 | 8 | 7 | | | |
| | | | 250 | 7 | 5 | | | |
| | | | 50 | 6 | 1 | | | |
| | | 6 | 5000 | 5 | 5 | 2 | 1 in 1211 (518-2829) | 0.083 (0.035-0.193) |
| | | | 1250 | 6 | 2 | | | |
| | | | 50 | 10 | 3 | | | |
| | | 24 | 5000 | 7 | 4 | 2 | 1 in 3371 (1614-7037) | 0.029 (0.014-0.062) |
| | | | 1250 | 8 | 4 | | | |
| | | | 50 | 8 | 1 | | | |
| | | 62 | 8000 | 4 | 3 | 2 | 1 in 3571 (1766-7223) | 0.028 (0.013-0.056) |
| | | | 5000 | 5 | 4 | | | |
| | | | 1250 | 7 | 3 | | | |
| | | | 50 | 7 | 0 | | | |
| **EXPER** | LT-HSCs | 0 | 5 | 14 | 6 | 2 | 1 in 14.8 (8.1 – 26.8) | 6.7 (3.7–12.3) |
| | | | 25 | 10 | 7 | | | |
| | | | 100 | 7 | 7 | | | |
| | | 6 | 5 | 8 | 4 | 2 | 1 in 15.2 (6.7 – 34.2) | 6.6 (2.9-14.8) |
| | | | 25 | 5 | 3 | | | |
| | | | 100 | 6 | 6 | | | |
| | | 24 | 10 | 8 | 0 | 3 | 1 in 80.6 (43.3 – 149.9) | 1.2 (0.06-2.3) |
| | | | 50 | 11 | 5 | | | |
| | | | 300 | 12 | 12 | | | |
| | | 72 | 50 | 13 | 4 | 5 | 1 in 293.7 (184.4 – 467.7) | 0.34 (0.12-0.54) |
| | | | 300 | 12 | 9 | | | |
| | | | 700 | 15 | 12 | | | |

**Table S1: Engraftment data for Limiting Dilution Assays reported in this study.**
Number of mice injected and number of mice engrafted for each of the doses tested in the limiting dilution experiments. Mice were considered engrafted if human cells ($hCD45^{++}$ &$GlyA^+$) >0.01% of Singlets and if >20 cells were present in any lineage determination gate. Engraftment measurements for all mice are reported in **Data S1**. LTRC: Long Term Repopulating Cell.

| Antibody | Manufacturer | Cat. # | Location used |
|---|---|---|---|
| Rat monoclonal anti-CD49f, PeCy5, clone GoH3 | BD | 551129 | Cambridge |
| Mouse monoclonal anti-CD38, PeCy7, clone HIT2 | Biolegend | 303516 | Cambridge |
| Mouse monoclonal anti-CD90, clone 5E10 | BD | 559869 | Cambridge |
| Mouse monoclonal anti- CD19, AlexaF700, clone HIB19 | Biolegend | 302226 | Cambridge |
| Mouse monoclonal anti-CD34, APCCy7, clone 581 | Biolegend | 343514 | Cambridge |
| Mouse monoclonal anti-CD45RA, BV421, clone HI100 | Biolegend | 304129 | Cambridge |
| Mouse monoclonal anti-GlyA, PE, clone HIR2 | BD | 340947 | Cambridge |
| Mouse monoclonal anti-CD45, PECy5, clone HI30 | Biolegend | 304010 | Cambridge |
| Mouse monoclonal anti-CD14, PECy7, clone M5E2 | Biolegend | 301814 | Cambridge |
| Mouse monoclonal anti-CD56, APC, clone HCD56 | Biolegend | 318310 | Cambridge |
| Mouse monoclonal CD11b, APCCy7, clone ICRF44 | Biolegend | 301342 | Cambridge |
| Mouse monoclonal anti-CD15, BV421, clone MC-48 | Biolegend | 125614 | Cambridge |
| Mouse monoclonal anti CD33, APC, clone P67.6 | BD | 345800 | Cambridge |
| Mouse monoclonal anti-CD3, APCCy7, clone HIT3a | Biolegend | 300318 | Cambridge |
| Mouse monoclonal anti-CD45, BV510, clone HI30 | Biolegend | 304036 | Cambridge |
| Rabbit monoclonal Phospho-Rb/Alexa Fluor 647 | Cell Signalling Technologies | 8974S | Cambridge |
| Mouse monoclonal anti-CD45, FITC, clone 2D1 | BD | 347463 | Toronto |
| Mouse monoclonal anti-CD14, PECy7, clone M5E2 | Beckmann Coulter | A07785 | Toronto |
| Mouse monoclonal anti-CD34, APCCy7, clone 581 | BD | custom made | Toronto |
| Mouse monoclonal anti-CD38, PeCy7, clone HB7 | BD | 335790 | Toronto |
| Mouse monoclonal anti-GlyA, PE, clone 11E4B-7-6 | Beckman Coulter | IM2211U | Toronto |
| Mouse monoclonal anti-CD14, PECy5.5, clone MΦP-9 | BD | B62209 | Toronto |
| Mouse monoclonal anti-CD33, APC, clone P67. | BD | 340474 | Toronto |
| Mouse monoclonal anti-CD19, AF700, clone HIB19 | Biolegend | 55792 | Toronto |
| Mouse monoclonal anti-CD3, APCCy7, clone SK7 | BD | 341090 | Toronto |
| Mouse monoclonal anti-CD45, BV450, clone HI30 | BD | 560367 | Toronto |
| Mouse monoclonal anti-CD45, VioBright B515, clone REA737 | Miltenyi | 130-130-165 | Toronto |

**Table S2: Antibodies used in this study**

This study used the antibodies in the table in the panels described below and referred to in the relevant methods sections:

**Panel A:** CD49f/PECy5 (clone GoH3, 1 in 100), CD38/PECy7 (clone HIT2, 1 in 100), CD90/APC (clone 5E10, 1 in 100), CD19/AlexaF700 (clone HIB19, 1 in 300), CD34/APCCy7 (clone 581, , 1 in 100) and CD45RA/BV421 (clone HI100, 1 in 100).
**Panel B:** GlyA/PE (clone HIR2, 1 in 1,000), CD45/PECy5 (clone HI30, 1 in 300), CD14/PECy7 (clone M5E2, 1 in 1000), CD56/APC (clone HCD56, 1 in 200), CD11b/APCCy7 (clone ICRF44, 1 in 300) and CD15/BV421 (clone MC-480, 1 in 200).
**Panel C:** GlyA/PE (clone HIR2, 1 in 1000), CD45/PECy5 (clone HI30, 1 in 300), CD14/PECy7 (clone M5E2, 1 in 1,000), CD33/APC (clone P67.6, 1 in 200), CD19/AlexaF700 (clone HIB19, 1 in 300), CD3/APCCy7 (clone HIT3a, 1 in 100) and CD45/BV510 (clone HI30, 1 in 500).
**Panel D:** CD45/FITC (clone 2D1, 1 in 100) CD45/PeCy5 (clone J33, 1 in 200), CD34/APCCy7 (clone 581, 1 in 200) and CD38/PeCy7 (clone HB7, 1 in 200).
**Panel E:** GlyA/PE (clone 11E4B-7-6), CD14/PeCy5-5 (clone MΦP-9), CD33/APC (clone P67.6, 1 in 100), CD19/AF700 (Clone HIB19), CD3/APCCy7 (clone SK7), CD45/BV450 (clone HI30), CD45/Vio Bright B515 (clone REA737) .
For Live/Dead discrimination Zombie Aqua (cat. no. 423101, 1 in 2,000) was included in Panel A and Sytox Blue**** (cat no. S34857, 1 in 2,000) was included in Panel D.

| Batch number | Time-point | Sorted | After QC |
|---|---|---|---|
| 1 | 0 h | 72 | 29 |
| | 6 h | 72 | 49 |
| | 24 h | 72 | 28 |
| | 72 h UNTR | 72 | 42 |
| | 72 h PD | 72 | 27 |
| 2 | 0 h | 94 | 56 |
| | 6 h | 94 | 85 |
| | 24 h UNTR | 95 | 58 |
| | 24 h PD | 95 | 80 |
| | 72 h UNTR | 95 | 82 |

**Table S3. Number of single cells before and after QC for Dataset 1** (EXPER conditions).

| Batch Number | Cell Type | Before QC | After QC | After outlier removal |
|---|---|---|---|---|
| Batch 1 | 0 h | 96 | 83 | 82 |
| | 62 h UNTR | 84 | 76 | 76 |
| | 62 h PD | 0 | 0 | 0 |
| Batch 2 | 0 h | 66 | 50 | 50 |
| | 62 h UNTR | 48 | 39 | 39 |
| | 62 h PD | 48 | 38 | 38 |
| Batch 3 | 0 h | 62 | 42 | 42 |
| | 62 h UNTR | 24 | 16 | 16 |
| | 62 h PD | 24 | 12 | 12 |
| Batch 4 | 0 h | 96 | 44 | 44 |
| | 62 h UNTR | 48 | 12 | 12 |
| | 62 h PD | 44 | 7 | 7 |

**Table S4. Number of single cells before and after QC for Dataset 2** (GT conditions).

| | Dataset 1 | | Dataset 2 | | | |
|---|---|---|---|---|---|---|
| **QC thresholds** | **Batch 1** | **Batch 2** | **Batch 1** | **Batch 2** | **Batch 3** | **Batch 4** |
| Number of Mapped reads (log10) | > 330,000 | > 190,000 | NA | NA | NA | NA |
| Number of nuclear genes (log10) | > 276,500 | > 180,000 | > $10^{5.2}$ | > $10^{5.2}$ | > 15,000 | > 100,000 |
| Ratio of genes to total number of reads | > 0.342 | > 0.2 | > 0.3 | > 0.3 | > 0.1 | > 0.2 |
| Number of genes with 10 reads per million | > 2100 | > 1000 | NA | > 1000 | > 1000 | > 1000 |
| Ratio of nuclear genes to number of mapped reads | > 0.818 | NA | NA | NA | NA | NA |
| Ratio of mitochondrial genes to number of mitochondrial and nuclear genes | NA | < 0.2 | < 0.2 | < 0.2 | < 0.2 | < 0.2 |
| Ratio of ERCC to number of mapped reads | NA | < 0.2 | < 0.4 | < 0.4 | < 0.2 | < 0.2 |
| Ratio of 'QC_no_feature' to total number of readsw | NA | < 0.36 | NA | NA | NA | NA |

**Table S5. Quality control thresholds (QC) for scRNA-seq experiments.**

QC thresholds used for filtering "good quality" single cells in each of the independent scRNAseq experiments. NA: not applied.

**Data S1. Experimental details and engraftment levels for mice used in primary transplantation experiments**

**Data S2. DESeq2 and GSVA output for the comparison of 0 h-early and 0 h-late LT-HSC subsets**

**Data S3. DESeq2 output for all pairwise comparisons of timepoints in the scRNAseq time course of LT-HSCs cultured in EXPER system**

The specific timepoints being compared are indicated in the name of each tab.

**Data S4. Lists of "ex vivo modulated genes" defined in this study**

Tab "ex_vivo_mod_genes_10010_EXPER" reports the union of all differentially expressed genes (by DESeq2 FDR<0.05) between any 2 pairwise comparisons in Integration 1 (0 h, 6 h, 24 h, 72 h; n=10,010 genes).
Tab "ex_vivo_mod_genes+PD_EXPER" reports the union of the 10,010 genes and of the genes differentially expressed between 0 h vs 72 h PD, 72 h PD vs 72 h UNTR, 0 vs 24 h PD and 24 h PD vs 24 h UNTR (n=10,903 genes).
Tab "ex_vivo_mod_genes+PD_GT" reports the union of all differentially expressed genes (by DESeq2 FDR<0.05) between any 2 pairwise comparisons in Dataset 2 (0 h, 62 h UNTR, 62 h PD; n=5,469 genes).

**Data S5. List of genes and GSVA pathways in each pattern identified by DEGpattern analysis**

**Data S6. List of maximally variable genes at each time point of scRNAseq timecourse, with associated Reactome pathways analysis**

These results pertain to the BASiCS variability analysis.
Tab "Maximally_variable_genes_time" reports the maximally variable genes corresponding to each timepoint. The other tabs show the full output of Reactome pathway analysis from these lists.

**Data S7. DESeq2 output and Reactome pathway analysis for comparisons of untreated and PD treated conditions**

Tabs "24hPD_vs24hUNTR_EXPER", "72hPD_vs72hUNTR_EXPER", and "62hPD_vs_62hUNTR_GT" tabs report the DEseq2 output for the analysis of differentially expressed genes between the conditions indicated in the tab name. The other tabs show the full output of Reactome pathway analysis from the differentially expressed genes (FDR < 0.05) in each of the lists.

All these supplementary data can be requested from the authors.
